# Periosteal skeletal stem cells can migrate into the bone marrow and support hematopoiesis after injury

**DOI:** 10.1101/2023.01.12.523842

**Authors:** Tony Marchand, Kemi E. Akinnola, Shoichiro Takeishi, Maria Maryanovich, Sandra Pinho, Julien Saint-Vanne, Alexander Birbrair, Thierry Lamy, Karin Tarte, Paul S. Frenette, Kira Gritsman

**Author notes:** These authors contributed equally to this work (Tony Marchand, Kemi E. Akinnola). Deceased (Paul S. Frenette).

## Abstract

Skeletal stem cells have been isolated from various tissues, including periosteum and bone marrow, where they exhibit key functions in bone biology and hematopoiesis, respectively. The role of periosteal skeletal stem cells in bone regeneration and healing has been extensively studied, but their ability to contribute to the bone marrow stroma is still under debate. In the present study, we characterized a whole bone transplantation model that mimics the initial bone marrow necrosis and fatty infiltration seen after injury. Using this model and a lineage tracing approach, we observed the migration of periosteal skeletal stem cells into the bone marrow after transplantation. Once in the bone marrow, periosteal skeletal stem cells are phenotypically and functionally reprogrammed into bone marrow mesenchymal stem cells that express high levels of hematopoietic stem cell niche factors such as Cxcl12 and Kitl. In addition, using *ex vivo* and *in vivo* approaches, we found that periosteal skeletal stem cells are more resistant to acute stress than bone marrow mesenchymal stem cells. These results highlight the plasticity of periosteal skeletal stem cells and their potential role in bone marrow regeneration after bone marrow injury.

## INTRODUCTION

Bone marrow mesenchymal stem cells (BM-MSCs) are rare self-renewing multipotent stromal cells which are capable of multilineage differentiation into osteoblasts, chondrocytes and adipocytes ^1–3^. BM-MSCs are mostly localized around the blood vessels and represent an important component of the hematopoietic stem cell (HSC) microenvironment, also referred to as the niche. BM-MSCs closely interact with HSCs and secrete factors, including the C-X-C motif chemokine ligand (CXCL12) and stem cell factor (SCF), that control their self-renewal, differentiation, and proliferation capacities ^4–9^. Several studies have used cell surface markers (CD51^+^, PDGFRα^+^, Sca-1^+^) or reporter mice (*Lepr*-cre, Nestin (*Nes*)-GFP, *Ng2*-cre) to describe distinct BM-MSC populations with significant overlap ^4–6,10–13^.

Recent studies have suggested that the periosteum, a thin layer of fibrous material that covers the surface of long bones, is a source of skeletal stem cells (SSCs) for bone regeneration ^14–18^. These periosteum SSCs (P-SSCs) have been described as sharing some characteristics with BM-MSCs ^14–16^. However, the relationship between these different stromal cell populations is poorly understood and the distinction between MSCs and SSCs remains controversial.

While the bone marrow is classically known as a major source of MSCs, the bone cortex represents a richer source of colony-forming units-fibroblasts (CFU-F) ^12,19,20^. It has been suggested that bone regeneration is mediated by both endochondral and intramembranous ossification and that the periosteum plays an important role in bone regeneration after injury^15,16,21–23^. However, there is little information on the function of P-SSCs, aside from their crucial role in bone healing and remodeling, and whether they contribute to bone marrow regeneration.

In the present study, we developed and characterized a whole bone transplantation model to study bone marrow regeneration, in which an intact adult femur is transplanted subcutaneously into a recipient mouse.^24^ Shortly after transplantation, bone marrow architecture in the transplanted femur, hereafter referred to as the graft, is severely altered with an expansion of adipocytes that mimics the fatty infiltration classically observed in the bone marrow after chemotherapy or radiation^25^. This initial destruction of the bone marrow microenvironment is followed by a progressive regeneration of blood vessels and the BM-MSCs network. The graft is progressively colonized by host-derived HSCs, allowing hematopoiesis to resume. Unexpectedly, we found that P-SSCs can migrate into the bone marrow and acquire BM-MSC niche functions, making them capable of supporting hematopoiesis through the *in vivo* expression of specific niche genes, such as *Cxcl12* and *Kitl*. In addition, we found that BM-MSCs and P-SSCs display different metabolic profiles, and that P-SSCs exhibit higher resistance to transplantation-induced stress. In conclusion, our study demonstrates the high plasticity of P-SSCs and highlights their potential contribution to bone marrow stroma regeneration after injury.

## RESULTS

### The whole bone transplant model recapitulates physiological regeneration of the bone marrow

In order to study the mechanisms involved in bone marrow regeneration, we employed a model system based on the subcutaneous transplantation of an intact adult femur into non-conditioned age and sex-matched recipient mice (**Figure 1A)**. The bone transplantation is followed by a rapid and massive depletion of bone marrow cells in the engrafted femur with a cell viability reaching below 10% at 24 hours after transplantation (**Figure S1A**). Notably, bone marrow necrosis following bone transplantation is associated with the replacement of hematopoietic cells in the graft femur by marrow adipocytes, similar to the effects of chemotherapy or irradiation ^25,26^ (**Figure S1B**). Following bone transplantation, we observed a progressive increase in bone marrow cellularity over time with no significant difference in cellularity between the graft and host femurs at five months following transplantation (**Figure 1B**). Regeneration of the bone marrow compartment is associated with a reduction in adipogenic infiltration, as revealed by staining with the anti-perilipin antibody at 4 months post-transplantation (**Figure S1B**). Additionally, the absolute number of graft BM-MSCs, defined as CD45^-^Ter119^-^CD31^-^CD51^+^CD140α^+^ cells by flow cytometry^12^, also increased over time, with no difference between the graft and host femurs at five months following transplantation (**Figure 1B**). We also observed a progressive increase in the number of Lin^-^Sca1^+^cKit^+^CD48^-^CD150^+^ phenotypic HSCs in the graft femur over time (**Figure 1B**). Although the absolute HSC numbers in the graft did not reach HSC numbers as detected in the host femur after five months (**Figure 1B**), no differences were observed in the numbers of hematopoietic progenitors or in the frequencies of hematopoietic cell populations between the host and graft femurs (**Figure S1C and S1D**). These data suggests that bone transplantation causes significant bone marrow injury, followed by a recovery process.

**Figure 1.**
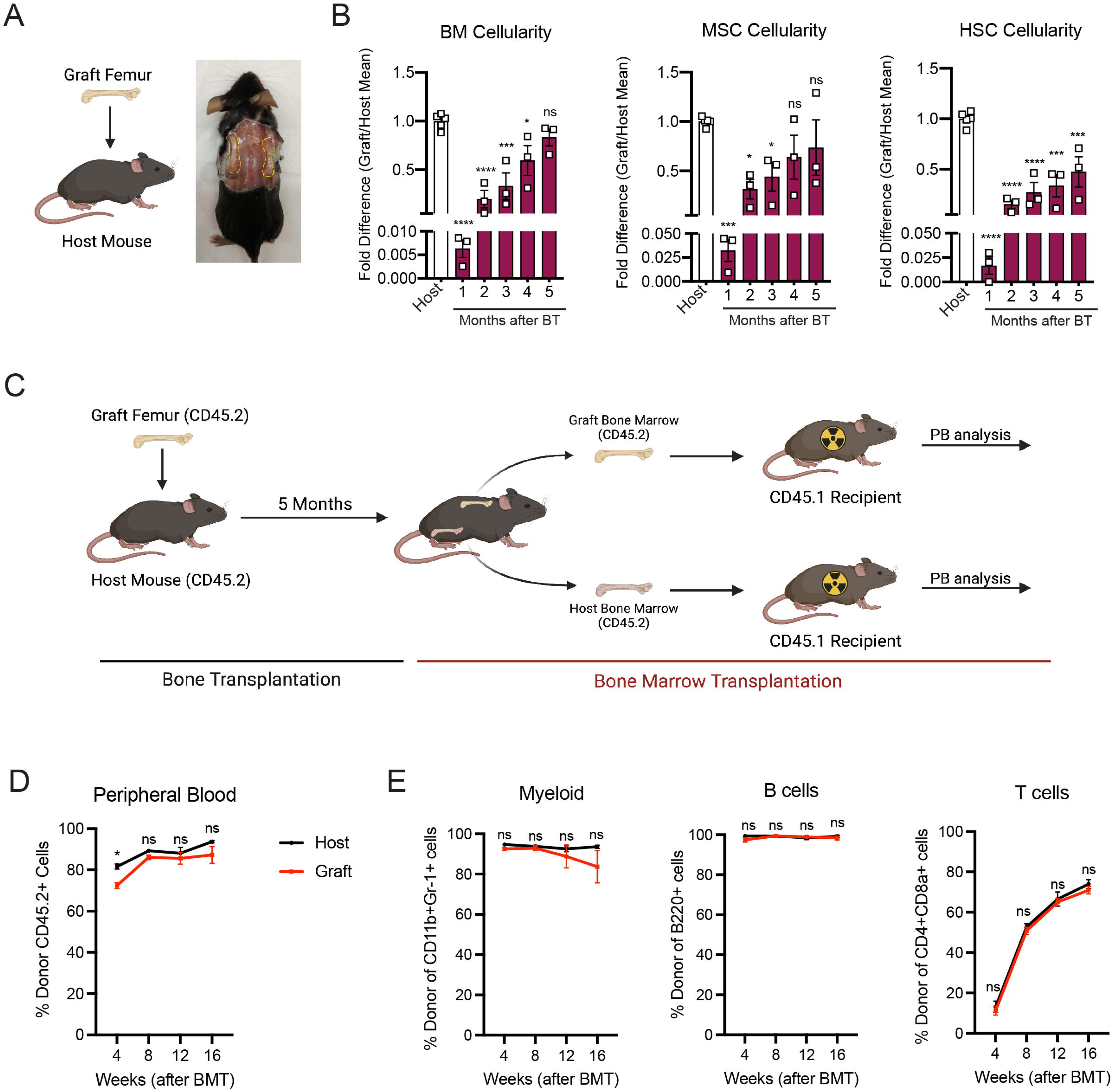
Whole bone transplantation is a good model to study bone marrow regeneration. A. Schematic and picture of the bone transplantation procedure. B. Fold difference quantification of graft femur/host femur cellularity normalized to mean host femur cellularity. Total graft bone marrow cells, BM-MSCs and HSCs were analyzed monthly until 5 months after bone transplantation (BT) (n=3). Ordinary one-way ANOVA with Dunnett multiple comparisons was used to determine statistical significance. C. Schematic illustration of the non-competitive repopulating assay after bone transplantation. D. Donor HSC contribution of graft and host recipients at 4 weeks after bone marrow transplantation (n=10). E. Quantification of tri-lineage (myeloid, B lymphoid, and T lymphoid cells) engraftment 4 weeks post transplantation (n=10).

Hematopoiesis is a highly regulated and essential process by which all the differentiated blood cells are produced. To investigate whether functional hematopoietic progenitors fully recover in graft femurs, we performed a non-competitive bone marrow transplantation assay, in which we transplanted either graft or host bone marrow cells into lethally irradiated recipients at five months after bone transplantation (**Figures 1C and S2A**). The survival of lethally irradiated recipients remained equal in both groups, with 100% of recipients surviving throughout the length of the experiment. Chimerism analysis revealed robust engraftment of recipient mice with hematopoietic cells derived from the engrafted femurs, indicating that HSCs and progenitors derived from graft femurs can sustain long-term hematopoiesis upon transplantation (**Figure 1D**). We also observed no significant differences in donor cell contribution to myeloid or lymphoid lineages between the two groups (**Figure 1E**). Altogether, these data establish our bone transplantation model as a useful tool to study bone marrow and HSC niche regeneration.

### Graft BM-MSCs are graft-derived and progressively express HSC niche factors during regeneration

Next, we aimed to determine the origin of the hematopoietic and stromal cell populations in the graft bone marrow. To achieve this, we took advantage of the Rosa^mT/mG^ and ubiquitin C (UBC) promoter driven-GFP mouse models, in which all cells are labeled by red and green reporters respectively ^27,28^. We transplanted femurs isolated from UBC-GFP mice into Rosa^mT/mG^ recipients and quantified endothelial cells, BM-MSCs, and hematopoietic cells in the graft (**Figures 2A and 2B**). Corroborating the results of Picoli et al., we observed that at five months post transplantation, over 98% of the graft BM-MSCs originated from the graft femur, while over 99% of the hematopoietic cells in the graft originated from the host mouse^24^ (**Figures 2C and 2D**). Interestingly, endothelial cells were derived from both the host and the graft, suggesting the contribution of different progenitors. To further assess endothelial regeneration within the context of bone transplantation, we conducted a transplantation experiment in which femurs from Cdh5 (VE-cadherin)-CreER; iTdTomato^29^ mice were transplanted into WT recipients. Confocal analysis was performed fifteen days (**Figure S3A**) and one month (**Figure S3B**) after transplantation. This revealed the presence of VE-cadherin-labeled vessels within the periosteum and crossing the cortical bone of the graft. This result demonstrates that endothelial regeneration occurs at an early stage following bone transplantation. Furthermore, when femurs from UBC-GFP mice were transplanted into Cdh5-CreER;iTdTomato mice, contributions of both recipient VE-cadherin-labelled vessels and UBC-GFP vessels were observed five months later. This result once again demonstrated the dual contribution of endothelial cells to the grafted femur (**Figure S3C**). Furthermore, we did not detect any cells derived from the graft in the host femurs (data not shown). These results are consistent with data obtained from ossicle-based experiments, where MSC-seeded ossicles are colonized by recipient-derived hematopoietic cells ^10,30–33^.

**Figure 2.**
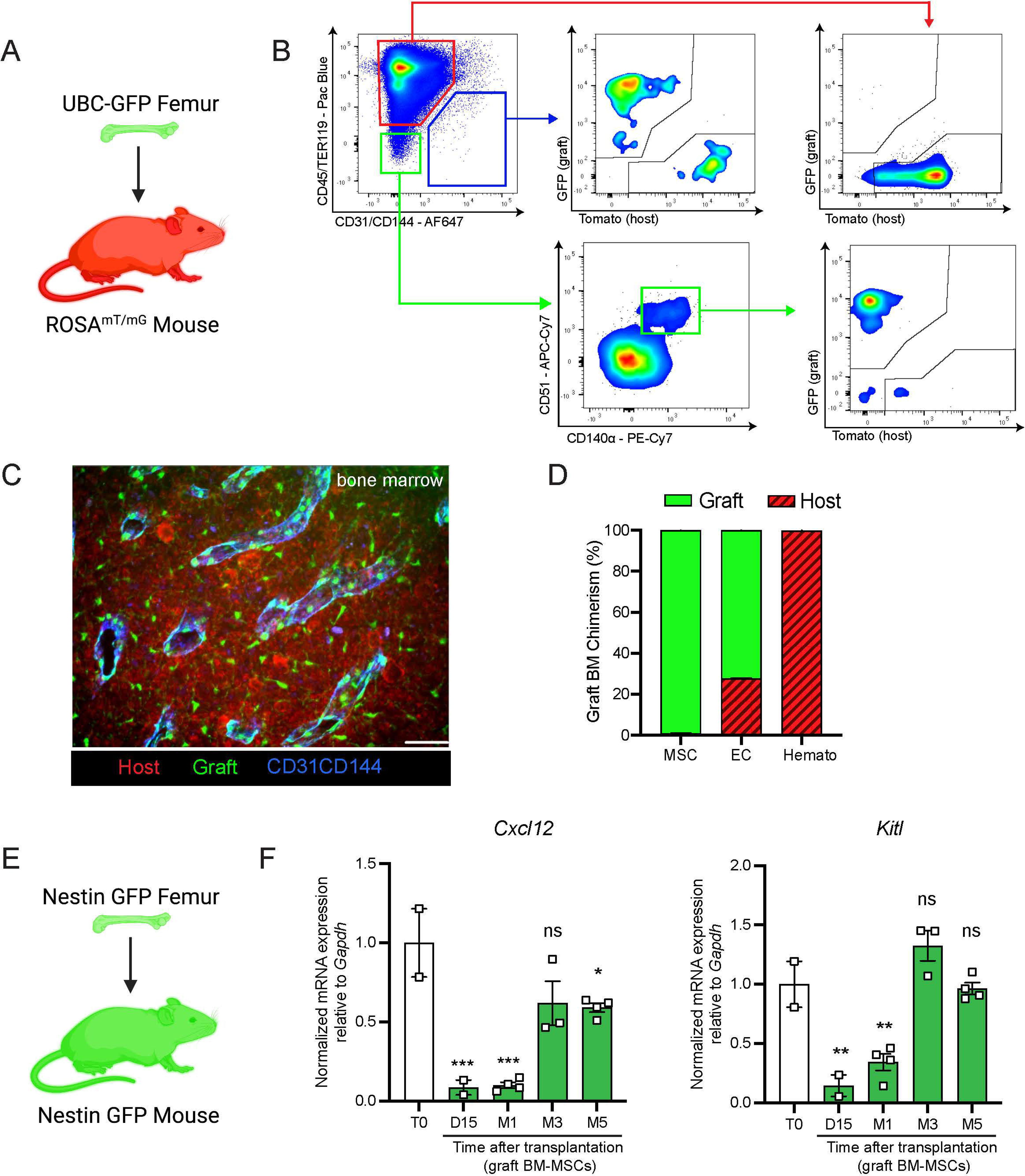
Regenerating BM-MSCs are graft-derived and express HSC niche factors. A. Schematic of a UBC-GFP femur transplanted into a Rosa^mT/mG^ mouse. B. Representative FACS plots showing the gating strategy to determine the origin of the different cell fractions in the graft 5 months after transplantation of a UBC-GFP femur into a Rosa^mT/mG^ mouse. C. Representative whole-mount confocal z-stack projections of a UBC-GFP bone transplanted into a Rosa^mT/mG^ recipient 5 months after transplantation. Vascularization was stained with anti-CD31 and anti-CD144 antibodies. Scale bar = 100µm (n=2 mice). D. Origin of graft BM-MSCs, endothelial cells (EC) and hematopoietic cells (Hemato) analyzed by flow cytometry 5 months after bone transplantation (n=2). E. Schematic of the *Nes*-GFP femur transplantation into a *Nes*-GFP mouse recipient. F. Quantitative RT-PCR analysis of mRNA expression of *Cxcl12* and *Kitl* expression relative to *Gapdh* in graft *Nes*-GFP^+^ BM-MSCs compared to steady-state *Nes*-GFP^+^ BM-MSCs at multiple time points after transplantation (n= 2-4 mice per time point). One-way ANOVA with Dunnett multiple comparisons was used to determine statistical significance. Data are represented as the mean ± SEM. Unless otherwise noted, statistical significance was determined using unpaired two-tailed Student’s t test. *p<0.05. ** p<0.01. *** p<0.001. ****p<0.0001.

BM-MSCs are a major constituent of the hematopoietic niche, secreting maintenance factors that support HSCs and hematopoietic progenitors^4–6,11,34,35^. To test for HSC niche supportive activity of graft BM-MSCs, we utilized *Nes*-GFP reporter mice to isolate BM-MSCs after bone transplantation. Previous work from our group has shown that *Nes*-GFP marks mouse MSCs with HSC-niche function within the bone marrow^10^. Since our previous analysis (**Figure 2D**) revealed that all of the graft BM-MSCs originate from the graft itself, we transplanted *Nes*-GFP femurs into *Nes*-GFP recipient mice and sorted CD45^-^Ter119^-^CD31^-^*Nes*-GFP^+^ BM-MSCs from donor and recipient mice for analysis at different time points (**Figure 2E**). Since we did not detect circulation of BM-MSCs between host and graft bone marrow in our bone transplantation experiments (data not shown), this strategy allowed us to compare host and graft BM-MSCs using equivalent markers. In addition to the previously described increase over time in the absolute number of BM-MSCs in the graft femur (**Figure 1B**), we observed a progressive increase in the expression of the HSC-niche genes *Cxcl12* and *Kitl* (which encodes Scf), reaching a plateau at five months post-transplantation (**Figure 2F**). Similarly, no differences were observed between host and graft femurs in the expression levels of additional niche factors, including *osteopontin* (*Opn)*^36,37^, *angiopoietin-1* (*Angpt1)*^38^, and *vascular cell adhesion molecule-1* (*Vcam1)*^39,40^ at five months (**Figure S2B**). In this experiment, host and graft BM-MSCs also had similar CFU-F activity at five months (**Figure S2C**). Altogether, these results show that by five months post-bone transplantation, the niche-supportive and *ex vivo* clonogenic functions of graft BM-MSCs are restored and are similar to the activity of native host BM-MSCs.

### P-SSCs but not BM-MSCs expand early after bone transplantation

SSCs are multipotent cells of the skeletal lineage that are important for bone development, repair, and homeostasis^41,42^. SSCs have been identified in the periosteum (P-SSCs), compact bone, and bone marrow^10,11,15,43^. Similar to BM-MSCs, P-SSCs have been shown to have CFU-F activity and the ability to differentiate into osteoblasts, chondrocytes, and adipocytes^14,16,44^. Due to the severe necrosis and depletion of the marrow cavity content that we observed following bone transplantation (**Figures 1B and S1A**), we hypothesized that cells derived from the compact bone and/or the periosteum could potentially contribute to stromal marrow regeneration. We first analyzed the cellularity of graft bone marrow, compact bone, and periosteum at different early time points following transplantation. While the number of live cells within the bone marrow and compact bone were drastically reduced in the first 24 hours post transplantation, we unexpectedly observed a significant but transient increase in live periosteal cells (**Figures 3A and S4A)**. Moreover, while most of the bone marrow cells were depleted shortly after transplantation (**Figures S1A and S4A**), cell viability was not affected in the periosteum within this time frame (**Figures 3A and S4B**). To quantify P-SSCs and BM-MSCs by flow cytometry, we used the combination of CD51 and CD200, as these markers have been well-validated in both tissues^14,16,45^. Within the CD45^-^Ter119^-^ CD31^-^ fraction of live periosteal cells, we confirmed that CD51^+^CD200^+^ P-SSCs had the highest CFU-F activity and were also capable of trilineage differentiation (**Figures S4C-S4E**). Flow cytometric analysis confirmed an expansion of P-SSCs starting at day 3 post transplantation and peaking at day 8 (**Figure 3B and S4F**). These results were confirmed by confocal microscopy analysis of *Nes*-GFP graft femurs transplanted into WT mice and stained for Periostin, a matricellular protein highly expressed by periosteal cells^15,46–49^. We detected an expansion of *Nes*-GFP^+^ skeletal progenitors^50^ within the periosteum with a peak at day 8 (**Figure 3C**), similar to the expansion kinetics that we detected by flow cytometry. Interestingly, by day 15, we could detect Nes-GFP^+^ cells outside of the periosteum layer stained by an anti-periostin antibody and in the compact bone (**Figure 3C**).

**Figure 3.**
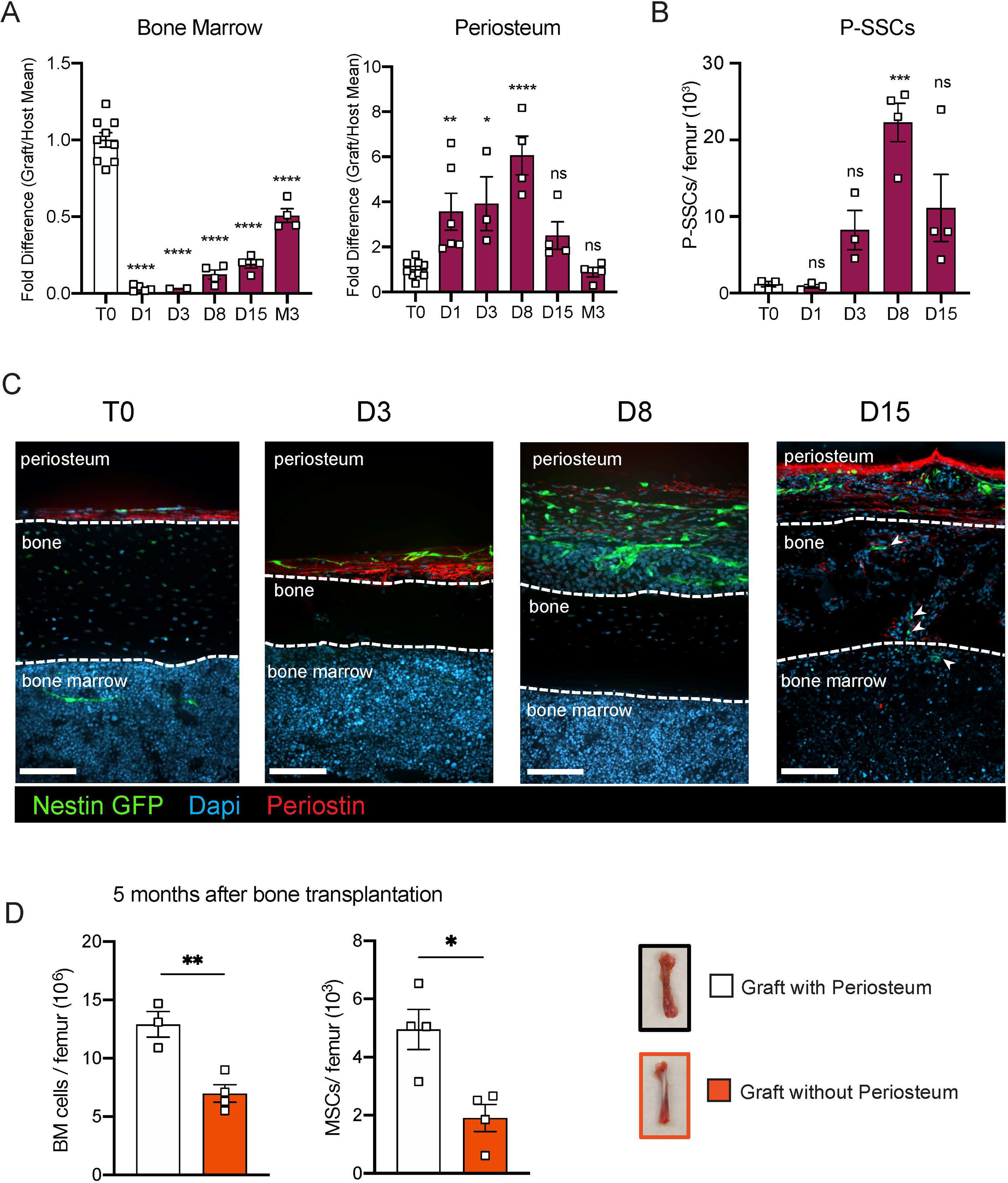
P-SSCs remain viable and expand after bone transplantation, in contrast to BM-MSCs. A. Flow cytometric quantification of fold difference of total graft bone marrow and periosteum cellularity to total steady state cellularity. Different time points early after transplantation were analyzed (n=2-8). One-way ANOVA with Dunnett multiple comparisons was used to determine statistical significance. B. Absolute number of CD45^-^Ter119^-^CD31^-^CD51^+^CD200^+^ P-SSCs at steady state and 1-, 8- and 15-days post transplantation (n=3-4 mice per time point). One-way ANOVA with Dunnett multiple comparisons was used to determine statistical significance. C. Representative whole-mount confocal z-stack projections of *Nes*-GFP^+^ bone graft at steady state, three-, eight-, and fifteen-days post transplantation. Three independent experiments yielded similar results. Arrowheads point at *Nes-*GFP^+^ cells within the bone and bone marrow. Scale bar = 100µm D. Total bone marrow cellularity and BM-MSC absolute number 5 months after transplantation of bones with or without intact periosteum (n=3-4 mice per group). Data are represented as the mean ± SEM. Unless otherwise noted, statistical significance was determined using unpaired two-tailed Student’s t test. *p<0.05. ** p<0.01. *** p<0.001. ****p<0.0001.

To evaluate the potential role of the periosteum in overall bone marrow regeneration, we compared the regenerative capacity of transplanted femurs with intact periosteum to that of femurs in which the periosteum was mechanically removed (**Figure 3D**). At five months after transplantation, total cellularity and BM-MSC number were significantly reduced in femurs lacking the periosteum, highlighting a potentially critical role for the periosteum and P-SSCs during bone marrow regeneration.

### P-SSCs are more resistant to stress than BM-MSCs

Due to the ability of P-SSCs to survive bone transplantation, as opposed to BM-MSCs, we explored the intrinsic differences between BM-MSCs and P-SSCs. Using RNA sequencing, we analyzed the transcriptional differences between CD51^+^CD200^+^ P-SSCs and BM-MSCs at steady-state. Gene set enrichment analysis (GSEA) revealed that P-SSCs were positively enriched for gene sets associated with stemness and negatively enriched for gene sets associated with proliferation (**Figure 4A**). These results are in line with qPCR analysis showing that P-SSCs express high levels of the cell cycle inhibitor genes *Cdkn1a* and *Cdkn1c,* and low levels of the cell cycle progression gene *Cdk4* at steady state **(Figures 4B**). Additionally, flow cytometric analysis revealed that, compared to BM-MSCs, P-SSCs are less metabolically active, as shown by decreased glucose uptake as assessed by 2-(N-(7-Nitrobenz-2-oxa-1,3-diazol-4-yl)Amino)-2-Deoxyglucose (2-NDBG) **(Figure 4C)**. This led us to hypothesize that P-SSCs are more resistant to stress than BM-MSCs.

**Figure 4.**
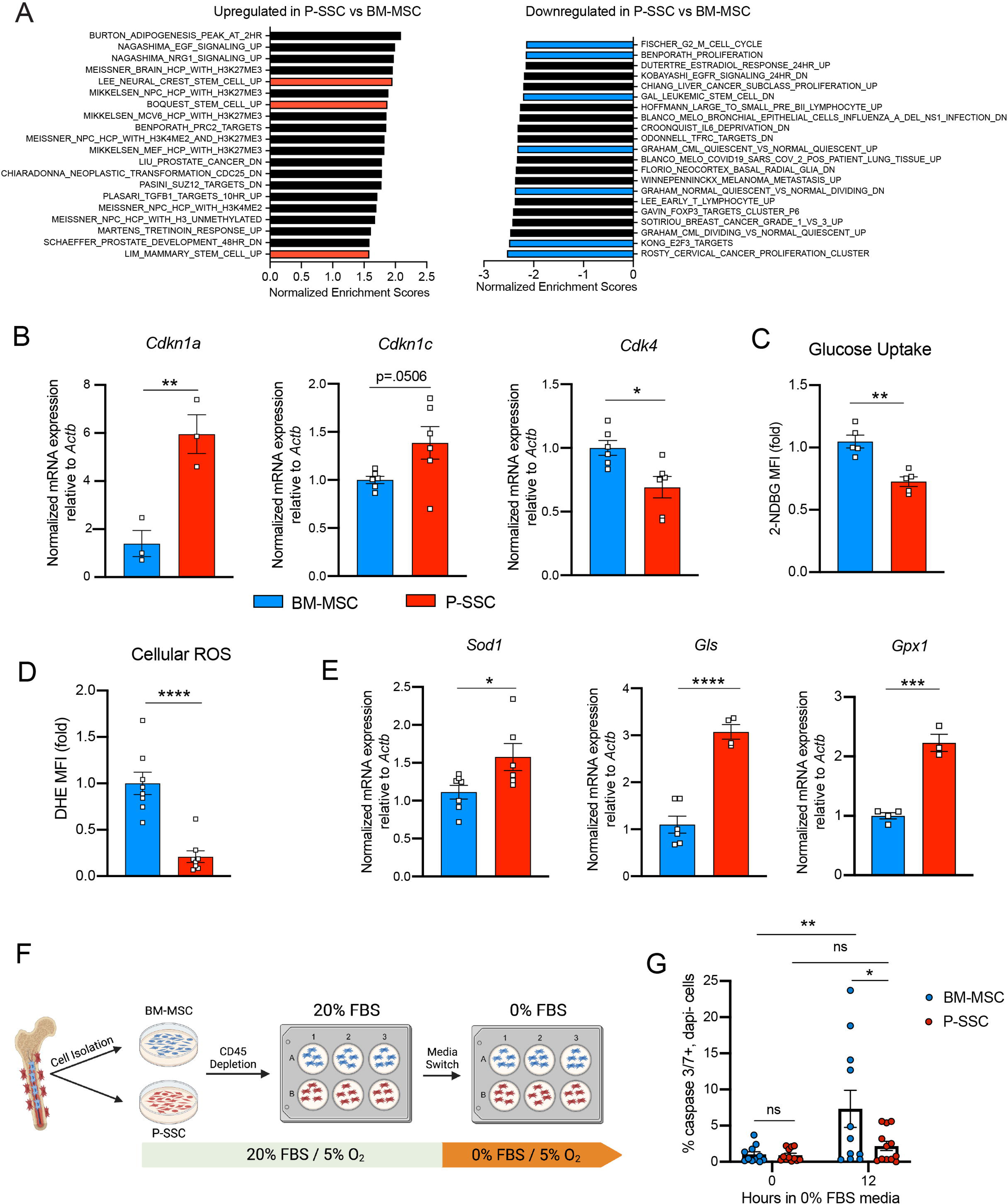
Periosteal SSCs have a metabolic profile conferring a resistance to stress. A. Gene set enrichment analysis (GSEA) plots comparing P-SSCs versus BM-MSCs at steady state (n=3 per group). B. Quantitative RT-PCR analysis of mRNA expression of *Cdkn1a*, *Cdkn1c*, *Cdk4* relative to *Actb* in sorted CD45^-^Ter119^-^CD31^-^CD51^+^CD200^+^ BM-MSCs and P-SSCs (n=3-6 per group). C. Flow cytometric analysis of glucose uptake at steady state in CD45^-^Ter119^-^CD31^-^CD51^+^CD200^+^ BM-MSCs and P-SSCs (n=5 per group). D. Quantification of cellular ROS at steady state in CD45^-^Ter119^-^CD31^-^CD51^+^CD200^+^ BM-MSCs and P-SSCs (n=8 per group). E. Quantitative RT-PCR analysis of mRNA expression of *Sod1*, *Gls* and *Gpx1* relative to *Actb* in sorted CD45^-^Ter119^-^CD31^-^CD51^+^CD200^+^ BM-MSCs and P-SSCs (n=3-7 per group). F. Schematic illustration of the protocol for the *in vitro* apoptosis assay. BM-MSCs and P-SSCs were isolated and digested before plating in a 10cm dish. At near confluence, cells underwent CD45 lineage depletion and plated into multi-well plates. At near confluence, medium was switched from 20% FBS to 0% FBS. Cells were analyzed at the time of medium switch and 12 hours. G. Percentage of apoptotic BM-MSCs and P-SSCs cultured under 5% O_2_ at baseline and 12 hours after being in 0% FBS serum conditions (n=11-12 per group). Two-way ANOVA with Tukey’s multiple comparisons test was used to determine statistical significance. Data are represented as the mean ± SEM. Unless otherwise noted, statistical significance was determined using unpaired two-tailed Student’s t test. *p<0.05. ** p<0.01. *** p<0.001. ****p<0.0001.

These observations led us to also use RNA sequencing to compare steady-state P-SSCs to P-SSCs at three days after bone transplantation. We chose the 72-hour time point to capture the time when P-SSCs are expanding but have not yet hit the peak of expansion at around day eight. We found that while steady-state P-SSCs and day three graft P-SSCs are quite distinct from steady-state BM-MSCs, there is a clear variance starting to occur between the two P-SSC populations **(Figure S5A)**. As early as three days post bone transplantation, we observed a downregulation of extracellular matrix (ECM) factors such as fibromodulin (*Fmod*) and nidogen 1 (*Nid1*) in the graft P-SCCs (**Figure S5B**). Additionally, gene set enrichment analysis (GSEA) comparing steady-state P-SSCs with graft P-SSCs, revealed an upregulation of cell cycle and DNA replication signatures and downregulation of ECM-receptor interaction and glutathione metabolism (**Figure S5C)**, indicating that a change in the P-SSCs phenotype occurs early after bone transplantation.

Low levels of reactive oxygen species (ROS) and high expression of antioxidant enzymes are mechanisms that help stem cells to avoid stress-induced cell death ^51,52^. Thus, we measured ROS levels by staining cells with the superoxide indicator dihydroethidium (DHE). At steady state, flow cytometric analysis revealed that P-SSCs had lower levels of cellular ROS than BM-MSCs (**Figure 4D**). Under physiological conditions, cells can maintain low ROS levels by expressing antioxidant enzymes. Indeed, qPCR analysis revealed higher expression of three major ROS-detoxifying enzymes genes: superoxide dismutase (*Sod1*), glutaminase (*Gls*) and glutathione peroxidase (*Gpx1*), in sorted CD51^+^CD200^+^ P-SSCs than in CD51^+^CD200^+^ BM-MSCs (**Figure 4E).** Altogether, these results suggest that P-SSCs are more stress-resistant than BM-MSCs, which may leave P-SSCs poised to proliferate in response to transplantation.

The location of the periosteum on the exterior of the bone, exposing P-SSCs to a higher oxygen tension than BM MSCs, may be a contributing factor to the observed differences in resilience between P-SSCs and BM-MSCs. Therefore, to determine whether the anatomic location of BM-MSCs and P-SSCs is the primary determinant of their differential stress response, we performed an *ex vivo* culture experiment designed to subject BM-MSCs and P-SSCs to equivalent levels of stress while in the same environment. After short-term *ex vivo* expansion of total bone marrow and periosteal cells, we performed a lineage depletion of CD45^+^ hematopoietic cells in both fractions. Purified P-SSCs and BM-MSCs were then maintained for 12 hours in serum-free culture media to stress the cells and mimic the nutrient deprivation that occurs immediately following bone transplantation **(Figure 4F)**. Flow cytometric analysis of activated caspase-3/7 revealed that after 12 hours in serum-free media, BM-MSCs exhibited a significantly higher level of apoptosis compared to P-SSCs (**Figures 4G and S5D**). These results suggest that P-SSCs are more intrinsically resilient than BM-MSCs, even when they are subjected to similar stress conditions in an equivalent environment.

### P-SSCs as a source of functional BM-MSCs during regeneration

As we observed that bone transplantation was followed by a depletion of bone marrow cellularity with an early expansion of P-SSCs, and that transplantation of bones without periosteum negatively impacts graft regeneration (**Figures 3D**), we hypothesized that proliferating P-SSCs migrate into the bone marrow and contribute to stromal regeneration. To test this hypothesis, we removed the periosteum from WT femurs and wrapped these femurs with the periosteum isolated from UBC-GFP mice (**Figure 5A and 5B**). We then transplanted these bones into WT host mice and observed GFP^+^ cells within the compact bone at five months after transplantation (**Figure 5C**), consistent with the well-described role of the periosteum in bone remodeling ^15,16,53^. Consistent with our hypothesis, we also observed GFP^+^ cells enwrapping endomucin-stained sinusoids and forming a network, similar to the perivascular nature of BM-MSCs^9,10^ (**Figure 5C**).

**Figure 5.**
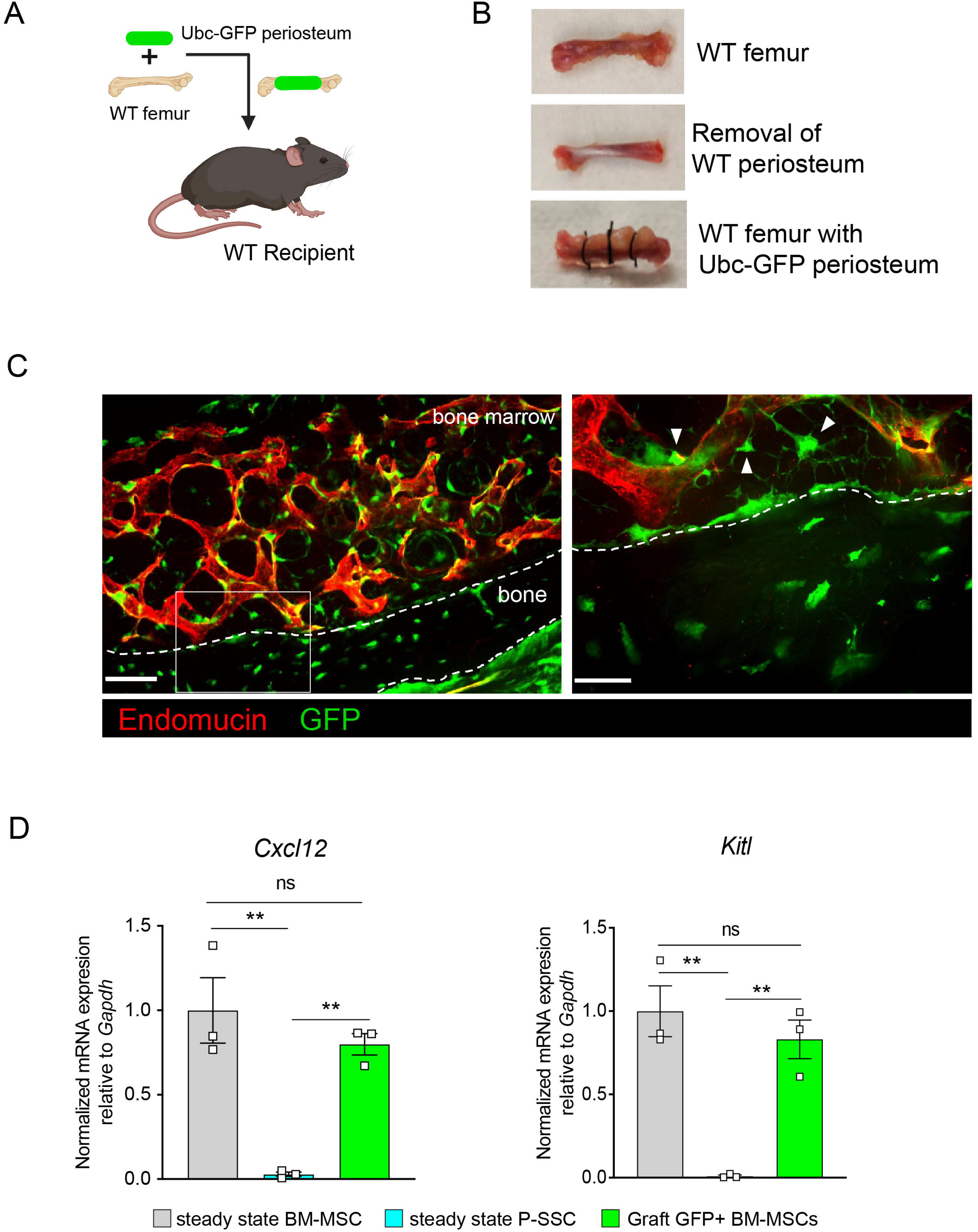
Periosteal SSCs migrate into the bone marrow and support stromal regeneration after bone transplantation. A. Schematic of the transplantation of a WT bone enwrapped with periosteum from a UBC-GFP mouse donor into a WT recipient mouse. B. Pictures illustrating the transplantation of a WT bone enwrapped with periosteum from a UBC-GFP mouse donor into a WT recipient mouse. C. Representative whole-mount confocal z-stack projections of wild-type bone graft enwrapped with periosteum from a UBC-GFP mouse donor into a WT recipient mouse 5 months after transplantation. Three independent experiments yielded similar results. Right panel: arrows pointing to GFP^+^ periosteum located perivascularly. Scale bar = 50µm (left panel) and 20µm (right panel) D. Quantification of *Cxcl12* and *Kitl* mRNA levels relative to *Gapdh* in sorted control CD45^-^Ter119^-^CD31^-^Nestin-GFP^+^ BM-MSCs, CD45^-^Ter119^-^CD31^-^CD51^+^CD200^+^ P-SSCs, and CD45^-^Ter119^-^CD31^-^CD51^+^CD200^+^GFP^+^ periosteum-derived graft BM-MSCs (n = 3-4 per group). One-way ANOVA with Tukey’s multiple comparisons was used to determine statistical significance. Data are represented as the mean ± SEM. Unless otherwise noted, statistical significance was determined using unpaired two-tailed Student’s t test. *p<0.05. ** p<0.01. *** p<0.001. ****p<0.0001.

Flow cytometric analysis of the graft at five months post-transplantation confirmed the presence of periosteum-derived GFP^+^ MSCs within the bone marrow cavity (**Figure S6A**). Importantly, while P-SSCs do not express *Cxcl12* or *Kitl* at steady state, periosteum-derived GFP+ BM-MSCs expressed these niche cytokines at a similar level to control sorted *Nes*-GFP+ BM-MSCs (**Figure 5D**). We also quantified the expression of the niche factors *Angpt1* and *Opn.* Similarly, we found that *Angpt1* expression in the graft GFP^+^ BM-MSCs reached the level of control BM-MSCs, while *Opn* expression was higher in GFP^+^ BM-MSCs than in control *Nes*-GFP+ BM-MSCs at five months after transplantation (**Figure S6B**). However, *Opn* has been shown to be upregulated in settings of inflammation, injury and migration^54–56^. Therefore, the moderate increase in *Opn* expression level in graft BM-MSCs compared to steady state BM-MSCs could be due to a residual inflammatory effect of bone transplantation. These results show that P-SSCs can both migrate into the bone marrow cavity and upregulate HSC maintenance genes to support hematopoiesis.

To confirm these results, we took advantage of a previously described transgenic mouse model in which an inducible Cre is placed under the promoter of the *Periostin* gene (*Postn^MCM^*), hereafter referred to as *Postn*-cre^ER^ ^57^. *Postn* encodes the secreted matricellular protein Periostin, and is highly expressed by periosteal cells and upregulated during bone healing and formation^15^. While *Postn* is expressed by multiple cell types, including but not limited to osteoblasts and fibroblasts^47,49,58^, its high expression in P-SSCs compared with BM-MSCs **(Figure S7A)** makes it a useful marker to distinguish endogenous BM-MSCs from periosteum-derived BM-MSCs after bone transplantation. We crossed the *Postn*-cre^ER^ line with ROSA26-loxP-stop-loxP-tdTomato reporter (Tomato) mice to be able to lineage trace P-SSCs. We transplanted femurs from *Postn*-cre^ER^;tdTomato mice into WT CD45.2 recipient mice, and then injected the host mice with tamoxifen shortly after transplantation to induce Cre recombination and Tomato expression in periosteal cells (**Figure 6A)**. At early time points following bone transplantation, we see Tomato^+^ cells confined to the periosteum and located perivascularly. At 21 days after transplantation, we observe Tomato+ cells migrating into the bone marrow (**Figure 6B**). By five months after transplantation, we observe robust Tomato^+^ labeling in the bone marrow located around the vasculature by confocal imaging **(Figure S7B)**, consistent with our prior observations **(Figure 5C).** Additionally, flow cytometric analysis revealed that an average of 85.4% (range: 65.0% – 94.5%) of the BM-MSCs within the engrafted bone marrow were Tomato^+^, indicating a periosteal origin (**Figures 6C and S7C**).

**Figure 6.**
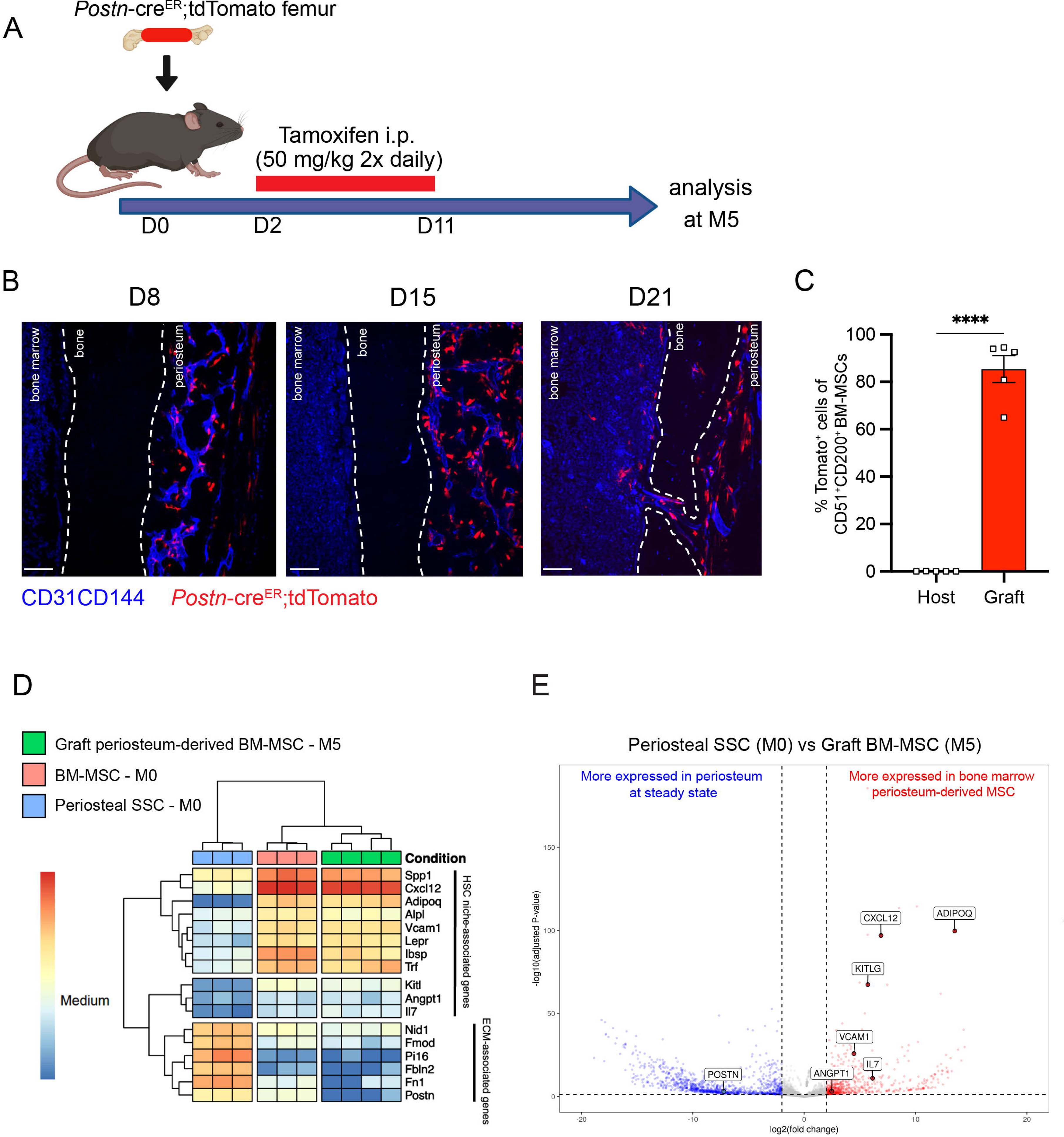
Periosteum-derived graft BM-MSCs adopt characteristics of baseline BM-MSCs, including the expression of HSC niche factors. A. Schematic illustration of the transplantation of a *Postn*-cre^ER^;tdTomato femur into a WT recipient mouse. B. Representative whole-mount confocal z-stack projections of transplanted *Postn*-cre^ER^;tdTomato femurs into a WT recipient 8-, 15-, and 21-days after transplantation. Two-three independent experiments yielded similar results. Scale bar = 100 µm C. Percentage of graft periosteum-derived BM-MSCs labeled Tomato^+^ five months after transplantation of a bone from a *Postn*-cre^ER^;tdTomato mouse into a WT recipient (n=5). D. Heat map expression level of selected genes defined by previous studies for HSC niche cells and extracellular matrix genes (n=3-4). E. Volcano plot of P-SSCs compared to graft BM-MSCs showing higher expression of HSC niche-associated genes in graft BM-MSCs. Data are represented as the mean ± SEM. Unless otherwise noted, statistical significance was determined using unpaired two-tailed Student’s t test. *p<0.05. ** p<0.01. *** p<0.001. ****p<0.0001.

We then wanted to assess our *Postn* mouse model using irradiation as a more physiologically relevant model of bone marrow injury, and to compare the effects of localized irradiation to femur transplantation on P-SSC plasticity. Single hindlimbs of *Postn-*cre^ER^;tdTomato mice were irradiated in order to compare the effects of irradiation within the same mouse (**Figure S7D**). While a clear difference in the architecture of bone marrow vasculature was observed between the non-irradiated and irradiated limbs, no difference in P-SSCs expansion or migration into the bone marrow was detected, contrasting with the findings in our femur transplant model (**Figure S7E**). This observation suggests that bone transplantation results in a more severe injury to the bone marrow compared to localized irradiation, thereby enabling the assessment of BM-MSCs’ recovery after more severe injury. However, it is also conceivable that stronger irradiation or a longer observation period would be necessary to observe a similar phenotype.

To examine changes in periosteum-derived BM-MSCs at the gene expression level, we performed bulk RNA sequencing on sorted CD51^+^CD200^+^Tomato+ BM-MSCs from graft *Postn*-cre^ER^;tdTomato femurs at five months after transplantation and on sorted steady-state CD51^+^CD200^+^ BM-MSCs and P-SSCs. Venn diagram and principal component analysis revealed that periosteum-derived graft BM-MSCs display a gene expression profile distinct from that of both steady state BM-MSCs and steady state P-SSCs (**Figures S8A and S8B**). Consistent with our previous tracing experiment using UBC-GFP periosteum **(Figure 5D)**, we observed an upregulation of HSC niche-associated maintenance genes in the five-month-old graft BM-MSCs compared to P-SSCs at steady state (**Figures 6D and 6E**). We also observed the downregulation of *Postn* and other extracellular matrix-related genes, such as fibronectin (*Fn1*) and fibromodulin (*Fmod*), in periosteum-derived graft BM-MSCs compared with P-SSCs at steady state (**Figure 6D and 6E**). Taken together, our results indicate that P-SSCs can be reprogrammed and adapt a niche-supportive phenotype akin to native BM-MSCs after migrating into the bone marrow following acute stress and subsequent regeneration.

## DISCUSSION

Bone marrow regeneration is a critical process that enables the recovery of hematopoiesis after injury such as irradiation or chemotherapy. Bone marrow mesenchymal stromal cells represent a key component of the bone marrow microenvironment. These stromal cells play a crucial role in regulating the self-renewal, differentiation, and proliferative properties of HSCs. The periosteum, a thin membrane that is highly vascularized and innervated, is located on the outside of the bone. This membrane contains numerous skeletal stromal cells, which play a pivotal role in maintaining the bone tissue and facilitating post-fracture healing. While the bone marrow microenvironment at steady state has been extensively studied, the mechanisms of bone marrow regeneration and stromal recovery remain poorly understood. Furthermore, data related to the functions of P-SSC beyond their established role in bone maintenance and fracture healing remain scarce. The objective of this study was to develop and characterize a model of whole bone transplantation in order to study bone marrow regeneration in mice. In this model, severe injury to hematopoietic and stromal cells within the bone marrow is induced, allowing for the investigation of the regeneration process of both cell populations. We found that the total graft bone marrow cellularity and BM-MSC cellularity demonstrate a gradual increase over time, ultimately reaching levels comparable to steady state levels by five months post-transplantation. The five-month time point was subsequently employed for further analyses. Prior research has emphasized the significance of stromal integrity for the recovery of HSCs following irradiation or chemotherapy ^59–61^. The findings of our study indicate that, while initially impacted by the stress associated with transplantation, BM-MSCs ultimately demonstrate their capacity to regenerate and sustain hematopoiesis within the engrafted femur. Furthermore, our findings indicate that hematopoietic progenitors derived from the graft femur are capable of engraftment in secondary recipient mice, thereby facilitating multi-lineage reconstitution. This model system recapitulates a recovering bone marrow microenvironment. Furthermore, it allows for genetic *in vivo* analysis of bone marrow regeneration. Consequently, bone transplantation can be employed as a valuable tool in future studies for investigating bone marrow regeneration.

The origin of endothelial progenitors in the bone marrow is not well defined. A recent study showed that during bone marrow regeneration after chemotherapy, sinusoidal and arteriolar vessels are derived from distinct progenitors^62^. Intriguingly, in the present study, we observed that endothelial cells within the graft originate from both the graft and host, which supports the hypothesis that different progenitors contribute to the bone marrow vascular network. Therefore, it is likely that endothelial regeneration in the graft, and vascular anastomoses between the host and graft femur are also important for the hematopoietic regeneration process. Further studies are needed to clarify the respective contributions of graft- and host-derived endothelial progenitors and the relative contributions of these different progenitor populations to regeneration of the vascular network and to hematopoietic regeneration.

Our results also highlight the high resilience and plasticity of P-SSCs and reveal their potential contribution to the bone marrow stromal network and bone marrow regeneration. At steady state, P-SSC do not express HSC maintenance genes, such as *Kitl* and *Cxcl12*, and the potential capacity of P-SSC to support HSCs has not been previously addressed. Unexpectedly, our results show that in our model, P-SSCs can migrate to the bone marrow and adopt a phenotype similar to that of BM-MSCs. Therefore, it is possible that P-SSCs can be harvested and manipulated as a source of BM-MSCs. Interestingly, we did not observe a similar migration and plasticity of P-SSCs in the context of localized femur irradiation. This observation suggests that femur transplantation introduces a greater stress on the bone marrow than irradiation, and that this increased level of stress is required to induce P-SSC plasticity and migration into the bone marrow.

In accordance with our findings, a recent study employing Gli1 to trace P-SSC demonstrated the localized expression of *Kitl* and *Cxcl12* by P-SSCs at the fracture site^43^. However, this model was not designed to specifically address the role of P-SSC in bone marrow regeneration. Furthermore, niche-specific genes were only expressed by cells adjacent to the fracture callus. Although we did not find a baseline difference in *Gli1* expression between BM-MSCs and P-SSCs in our RNA sequencing analysis, this may be due to the differences in surface markers and reporters used to identify P-SSCs and BM-MSCs. In the same study, authors did not find expression of reporters in the periosteum using the *Postn*-cre^ER^ mice. It is likely that this discrepancy is attributable to differences in generation of the Postn-Cre mice used. Indeed, previous studies have demonstrated that, compared to steady state, the *Postn* gene is upregulated following the activation of P-SSC, which is clearly evident in the context of whole bone transplantation.^63^ The results of our flow cytometry and imaging analyses demonstrate that P-SSC is specifically labelled at early time points following the transplantation of grafts from Postn-Cre^ER^ mice. Therefore, we show a novel application for the use of the inducible *Postn*-cre^ER^ mice to differentiate between BM-MSCs and P-SSCs *in vivo.* Accordingly, Duchamp et al. demonstrated that periostin contributes to the highly regenerative nature of P-SSCs compared to BM-MSCs^15^. While previous studies have used *Prx1* and *Ctsk-*Cre models, these models do not adequately allow the distinction between P-SSCs and BM-MSCs, likely due to their common embryonic origin^15,16,41^. Periostin is a well-studied protein that has been shown to interact with extracellular matrix proteins and plays a key role in tissue regeneration and cancer progression, promoting proliferation, invasion, and anti-apoptotic signaling ^46,64–66^. Therefore, it is possible that periostin contributes to P-SSC proliferation and migration into the bone marrow, which would be an interesting area for future investigation.

Additionally, our findings illustrate the differential stress response between BM-MSCs and P-SSCs. While BM-MSCs are renowned for their resilience to stress, our findings illustrate that P-SSCs exhibit an even greater resistance to stress, which is attributed, at least in part, to their distinctive metabolic profile^13,67^. Given that differences in apoptosis were observed between BM-MSCs and P-SSCs, even when they were cultured ex vivo under identical stress culture conditions with identical oxygen tension, it can be concluded that the observed differences between P-SSCs and BM-MSCs are due to intrinsic cellular properties rather than their anatomical location. Nevertheless, additional studies are required to elucidate the underlying mechanism responsible for the observed relative stress resistance of P-SSCs.

In conclusion, we have used a whole bone transplantation model to study bone marrow regeneration *in vivo* in response to acute injury using genetic tools. Our study has shown that through their high level of resilience and plasticity, P-SSCs can facilitate BM-MSC regeneration. Our data suggest that P-SSCs are able to contribute to hematopoietic cell recovery under stress conditions.

## EXPERIMENTAL PROCEDURES

### Mice

Mice were maintained under specific pathogen-free conditions in a barrier facility in microisolator cages. This study complied with all ethical regulations involving experiments with mice, and the Institutional Animal Care and Use Committee of Albert Einstein College of Medicine approved all experimental procedures, based on protocol #00001101. C57BL/6J mice were bred in our facilities or ordered from Jackson Laboratory. B6.129-Postn^tm2.1^(cre/Esr1*)^Jmol^/J^57^ were ordered from Jackson Laboratory and then bred in our facilities. Nestin-GFP, *Gt(ROSA)26Sor^tm^*^4^*^(ACTB-tdTomato,-EGFP)Luo^/J* (Rosa^mT/mG^), C57BL/6-Tg(UBC-GFP)30Scha/J mice were bred in our facilities, Cdh5-CreER; iTdTomato mice were kindly provided by R. H. Adams (Max Planck Institute for Molecular Biomedicine, Germany) and then bred in our facilities. Unless otherwise specified, 6-to-12-week-old mice were used for the experiments. For all analytical and therapeutic experiments, sex-matched animals from the same age group were randomly assigned to experimental groups.

### Bone transplantation procedure

Donor mice were anesthetized with isoflurane and euthanized by cervical dislocation. Femurs were isolated and preserved in an ice-cold phosphate-buffered saline (PBS) solution with 1% fetal bovine serum (FBS). Recipient mice were anesthetized with a ketamine/xylazine intraperitoneal injection (10 µL/g). Donor femurs were subcutaneously implanted in the back of the recipient mice, and the skin was sutured with a non-absorbable polyamide 5/0 silk. Mice were allowed to recover under a heat lamp until awake and monitored daily for up to a week post-surgery.

### *In vivo* treatment

For lineage tracing experiments using femurs from Postn^tm2.1^(cre/Esr1*)^Jmol^/J donor mice, tamoxifen (1mg/mouse) was administered intraperitoneally to recipient mice twice daily for 10 consecutive days starting at day 2 post-transplantation.

### Bone marrow transplantation

Non-competitive repopulation assays were performed using CD45.1 and CD45.2 mice. Recipient mice were lethally irradiated (12 Gy, two split doses) in a Cesium Mark 1 irradiator (JL Shepherd & Associates). A total of 1 x 10^6^ CD45.2^+^ bone marrow nuclear cells from either the graft or host femurs were obtained at five months after transplantation and injected retro-orbitally into irradiated CD45.1^+^ mice. Mice were bled retro-orbitally every 4 weeks after bone marrow transplantation, and peripheral blood was analyzed for engraftment and repopulation up to 16 weeks.

### Preparation of single cell suspensions

To isolate P-SSCs, muscle tissue was carefully removed using scissors and intact bones were submerged for 30 minutes in ice-cold PBS with 1% fetal bovine serum (FBS). The periosteum was carefully removed with a surgical blade, and mechanical dissociation was performed using scissors. Enzymatic dissociation was performed by incubating the periosteum fragments for 45 minutes at 37°C in digestion buffer (Hank’s balanced salt solution (HBSS, Gibco) containing 1 mg.ml^−1^ collagenase type IV (Gibco) and 2 mg.ml^−1^ dispase (Gibco)) on a rotator. Bone marrow cells were obtained by flushing and dissociating using a 1-ml syringe with PBS via a 21-gauge needle. For analysis of stromal and endothelial cell populations, intact bone marrow plugs were flushed into digestion buffer using 21- or 25-gauge needles and incubated at 37 °C for 30 min with manual mixing every 10 mins. After bone marrow and periosteum isolation, the remaining compact bone was crushed, mechanically dissociated using scissors as previously described ^45^ and digested in the digestion buffer, rotating for 45 minutes at 37°C. Enzymatic digestion was stopped by adding ice-cold PEB buffer (PBS with 0.5% BSA and 2mM EDTA).

### Flow cytometry and cell sorting

For FACS analysis and sorting, red blood cells were lysed (distilled H_2_O containing 155mM ammonium chloride, 10mM potassium bicarbonate and 0.5M EDTA) and washed in ice-cold PEB (PBS containing 0.5% BSA and 2 mM EDTA) before staining with antibodies in PEB for 20 minutes on ice. Dead cells and debris were excluded by FSC (forward scatter), SSC (side scatter) and DAPI (4’,6-diamino-2-phenylindole; Sigma). FACS analyses were carried out using BD LSRII flow cytometry (BD Biosciences) and cell sorting experiments were performed using a MoFlo Astrios (Beckman Coulter). Data were analyzed with FlowJo 10.4.0 (LCC) and FACS Diva 6.1 software (BD Biosciences). Antibodies used for FACS can be found in Supplementary table 1. For metabolic assays, cells were first stained with cell surface markers prior to labeling with metabolic dyes. For cellular ROS quantification, cells were incubated with dihydroethidium (5uM; Molecular Probes) for 20 minutes at 37°C in PBS. Glucose uptake quantification was performed by incubating the cells in DMEM without glucose (Gibco) containing Glutamax (1:100; Gibco) and 2-NBDG (17µmol mL^-1^; Cayman Chemical Company) for 30 minutes at 37°C.

### CFU-F assays

For CFU-F and stromal cell culture, CD45^-^Ter119^-^CD31^-^CD51^+^CD200^+^ stromal cells isolated from bone marrow and periosteum were sorted and plated at a clonal density (1,000 cell/well) in α-MEM (Gibco) containing 20% FBS (HyClone), 10% MesenCult Stimulatory supplement (StemCell Technologies) and 1% Penicillin-Streptomycin. Half of the medium was changed at day 7. Cells were cultured for 12-14 days, at the end of which the colonies were scored.

### Osteogenic, adipogenic and chondrogenic differentiation assays

Trilineage differentiation assays towards the osteogenic, adipogenic, and chondrogenic lineages were performed as previously described^12^, with minor modifications. Briefly, cells were treated with StemXVivo Osteogenic, Adipogenic, or Chondrogenic mouse differentiation media, according to the manufacturer’s instructions (R&D Systems). All cultures were maintained with 5% CO2 in a water-jacketed incubator at 37°C. Osteogenic differentiation was revealed by Alizarin Red S staining. Adipocytes were identified by the typical production of lipid droplets and Bodipy (Invitrogen) staining. Chondrocytes were revealed by Alcian Blue staining.

### *Ex vivo* culture nutrient deprivation assay

Whole bone marrow from 1 femur and whole periosteum from 2 femurs were isolated and digested as previously described and plated in α-MEM (Gibco) containing 20% FBS (HyClone), 1% penicillin-streptomycin, 1% L-glutamine and βFGF. The medium was changed every 3-4 days. Once a plate reached near confluence, CD45 lineage depletion was performed on both bone marrow and periosteum fractions. Cells were then counted and plated in 12- or 24-well plates at approximately 5000 cells/cm^2^. Once the plates reached near confluence, media was switched to α-MEM without FBS, 1% L-glutamine and βFGF. 12 hours after the medium was switched to 0% FBS medium, the cells were trypsinized, spun down, and stained for cell surface markers. After 4-5 days, flow cytometric apoptosis quantification was performed using the CellEvent Caspase 3/7 kit (ThermoFisher) following the manufacturer’s recommendations.

### Immunofluorescence imaging of bone sections

To stain blood vessels, anti-CD31 and anti-CD144 antibodies were injected intravenously into mice (10 μg, 20 μL of 0.5 μg.μL^−1^) and mice were sacrificed for analysis at 10 min after injection. For frozen sections of long bones, femurs and tibias were fixed in 4% paraformaldehyde (PFA) overnight at 4 °C. For cryopreservation, the bones were incubated sequentially in 10%, 20%, and 30% sucrose/PBS at 4 °C for 1h each and embedded and flash frozen in SCEM embedding medium (SECTION-LAB). Frozen sections were prepared at 20μm thickness with a cryostat (CM3050, Leica) using the Kawamoto’s tape transfer method ^68^. For immunofluorescence staining, sections were rinsed with PBS, post-fixed with 4% cold PFA for 10 min, followed by blocking with 20% donkey serum (DS; Sigma) in 0.5% Triton X-100/PBS for 3 h at room temperature (20–25 °C). For perilipin staining, sections were incubated for 1 hour at room temperature in saturation buffer (PBS-donkey serum 10%). The rabbit polyclonal anti-perilipin antibody (clone: D1D8; Cat: 9349; Cell Signaling Technology) was used at 1:100 dilution in 2% Donkey serum 0.1% Triton X-100/PBS overnight at 4 °C. Periostin staining was performed using whole mount femur imaging. The bone marrow was exposed by shaving the bone using a cryostat (CM3050, Leica). Shaved femurs were fixed 30 minutes at 4°C in PBS/PFA 4%. Samples were then incubated in the saturation buffer (PBS-donkey serum 10%) during 1 hour at room temperature. Polyclonal goat anti-periostin antibody (Cat: AF2955; R&D) and monoclonal rat anti-endomucin antibodies (clone: V.7C7; Cat: sc-65495; Santa Cruz) were used at a 1:100 dilution overnight at 4°C in PBS-donkey serum 2%. When necessary, primary antibody staining was followed by 3 washes with 2% DS 0.1% Triton X-100/PBS and a 30 min incubation with Alexa Fluor 568 or Alexa Fluor 488-conjugated secondary antibodies (Invitrogen) and 0.2% DAPI (4′, 6-diamino-2-phenylindole; Sigma).

### Image acquisition

All images were acquired at room temperature using a Zeiss Axio examiner D1 microscope (Zeiss) with a confocal scanner unit (Yokogawa) and reconstructed in three dimensions with Slide Book software (Intelligent Imaging Innovations). Image analysis was performed using both Slide Book software (Intelligent Imaging Innovations) and the Fiji build of ImageJ (NIH).

### Single limb irradiation

Mice were anesthetized using isoflurane chamber and kept anesthetized throughout the procedure. Mice were irradiated using the Small Animal Radiation Research Platform, SARRP (XStrahl, Surrey, UK). Whole body lead shielding protected the mice except for one hindlimb which was protruding out from under the shield and irradiated to 20 Gy for 319 seconds in a single fraction. Mice were left under the heat lamp until they recovered.

### RNA isolation and quantitative real-time PCR (q-PCR)

mRNA was purified using the Dynabeads® mRNA DIRECT™ Micro Kit (Life technologies - Invitrogen) by directly sorting stromal cells into lysis buffer, and reverse transcription was performed using RNA to cDNA EcoDry^TM^ Premix (Clontech – Takara Bio) following the manufacturer’s instructions. The SYBR green (Roche) method was used for quantitative PCR using the QuantStudio 6 Flex system (Applied Biosystems, ThermoFisher). All mRNA expression levels were calculated relative to *Gapdh* or *Actb*. Supplementary table 2 lists the primer sequences used.

### RNA sequencing and analysis

Total RNA from 1000-3000 sorted steady BM-MSCs, steady state P-SSCs, graft P-SSCs, and graft BM-MSCs was extracted using the RNAeasy Plus Micro kit (Qiagen) and assessed for integrity and purity using an Agilent Bioanalyzer. When applicable, RNA from two mice was combined; however, each replicate contained RNA from distinct mice. RNA-seq data generated from Illumina Novaseq6000 were processed using the following pipeline. Briefly, clean reads were mapped to the mouse genome (GRCm38) using Spliced Transcripts Alignment to a Reference (STAR 2.6.1d). Gene expression levels were calculated and differentially expressed genes were identified using DESeq2 and enriched using clusterProfiler. All RNA sequencing data are available under the SuperSeries dataset GSE222272 in GEO omnibus.

### Statistical analysis

All data are presented as the mean±S.E.M. N represents the number of mice in each experiment, as detailed in the figure legends. No statistical method was used to predetermine sample sizes; sample sizes were determined by previous experience with similar models of hematopoiesis, as shown in previous experiments performed in our laboratory. Statistical significance was determined by an unpaired, two-tailed Student’s t-test to compare two groups or a one-way ANOVA with multiple group comparisons. Statistical analyses were performed, and data presented using GraphPad Prism 8 (GraphPad Software), FACS Diva 6.1 software (BD Biosciences, FlowJo 10.4.0 (LLC), Slide Book Software 6.0 (Intelligent Imaging Innovations) and QuantStudio 6 Real-Time PCR Software (Applied Biosystem, Thermo Fisher). *P<0.05, **P<0.01, ***P<0.001, ****P<0.0001.

## DATA AVAILABILITY

RNA sequencing data from this study are available at accession number GSE222272 in the GEO Omnibus.

## ACKNOWLEDGMENTS

We would like to thank Colette Prophete and Daqian Sun for technical assistance and Lydia Tesfa and the Einstein Flow Cytometry Core Facility for expert cell sort assistance. We thank Charles Brottier and Robert Dubin of the Albert Einstein College of Medicine Computational Biology core facility for their help with analysis of RNA sequencing data. This work was supported by the National Institutes of Health (NIH) Grant 5R01DK056638 (to P.S.F. and K.G.), administrative supplement R01DK056638-23S1 (to K.E.A.), R01DK112976 (to P.S.F.), R56DK130895 (to K.G.), R01DK130895 (to K.G.), R01HL162584 (to S.P.), the NIH training Grant T32GM007288-50 (to K.E.A), and NYSTEM IIRP C029570A (to P.S.F.). T.M. was supported by the Fondation ARC pour la Recherche sur le Cancer, the Association pour le Développement de l’Hématologie Oncologie, the Société Française d’Hématologie, the Centre Hospitalier Universitaire de Rennes, and the Philip Foundation. S.T. was supported by the Japan Society for the Promotion of Science (JSPS) Postdoctoral Fellowship for Research Abroad, the Uehara Memorial Foundation Research Fellowship, and the NYSTEM Empire State Institutional Program in Stem Cell Research. M.M. was supported by the EMBO European Commission FP7 (Marie Curie Actions; EMBOCOFUND2012, GA-2012-600394, ALTF 447-2014), by the New York Stem Cell Foundation (NYSCF) Druckenmiller fellowship, and by the American Society of Hematology (ASH) Research Restart Award. S.P. was also supported by a Longevity Impetus Grant from Norn Group. This work was also supported by the Albert Einstein Cancer Center core support grant P30CA013330. Experimental figure illustrations were created using BioRender. The content is solely the responsibility of the authors and does not necessarily represent the official views of the National Institutes of Health.

## AUTHOR CONTRIBUTIONS

T.M, K.E.A, P.S.F, and K.G. designed the experiments. T.M, K.E.A, S.T., M.M. and S.P. performed the experiments and analyzed the data. T.M. and K.A. prepared the figures, and T.M., K.E.A., M.M., S.P., S.T. and K.G. wrote and edited the manuscript. P.S.F., K.G., T.L., and K.T. acquired funding for the study. J.S.V. performed the GSE222271 data analysis. A.B. conceptualized the whole bone transplantation procedure. All authors have read and approved the submitted manuscript.

## DECLARATION OF INTERESTS

The authors declare no competing interests.

## Notes

https://www.ncbi.nlm.nih.gov/geo/query/acc.cgi?acc=GSE222272

### Competing Interest Statement

The authors have declared no competing interest.

### Summary of Updates

New data has been added to compare the femur transplant model to a localized bone irradiation model and to visualize the re-endothelialization of the femur grafts. In addition, new RNA sequencing data has been added from periosteal SSCs from grafted femurs at an earlier time point to examine earlier changes in periosteal cells upon transplantation.

