## Supplementary Data for "Periosteal skeletal stem cells can migrate into the bone marrow and support hematopoiesis after injury"

Supplemental Figure 1

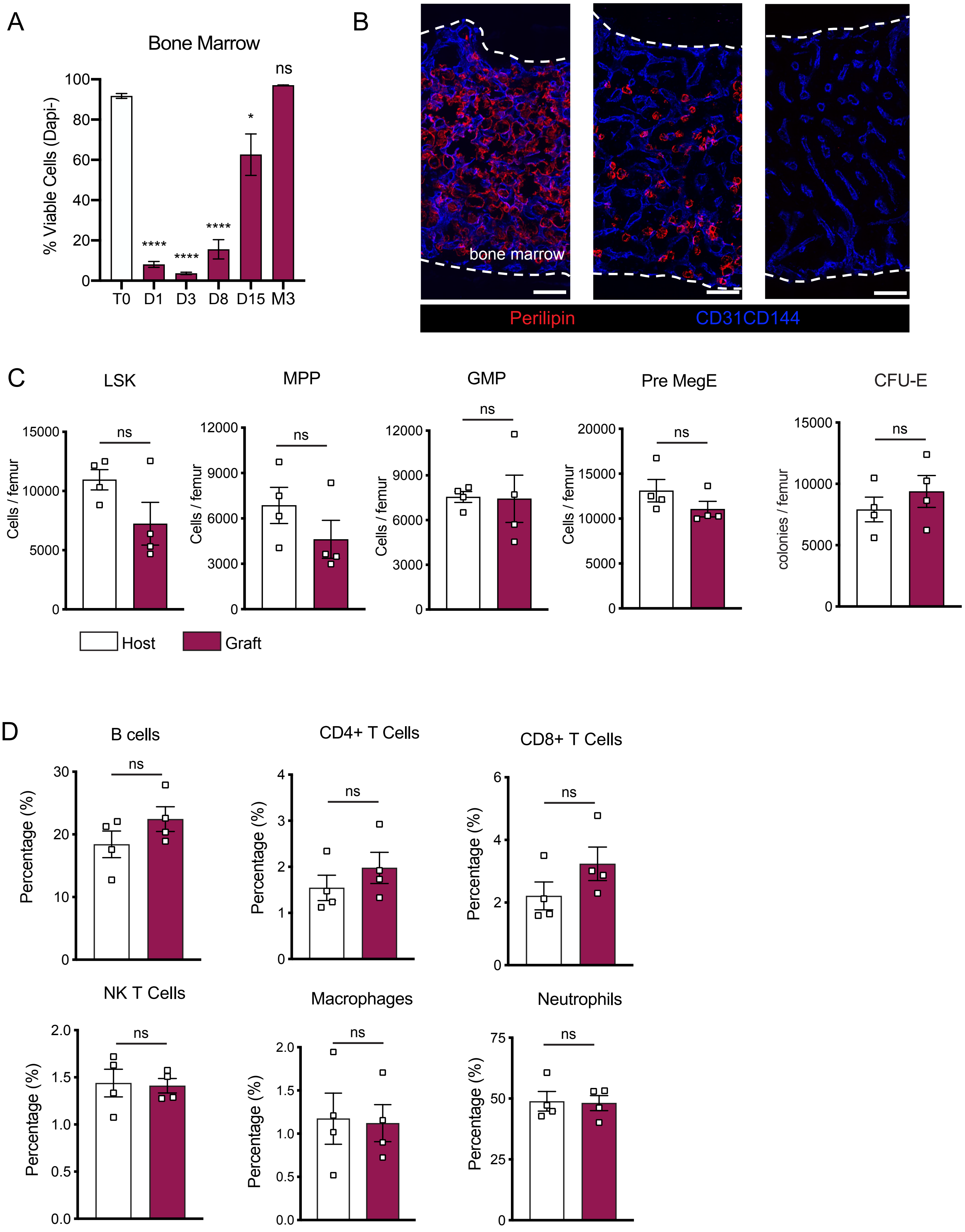

**Supplemental Figure 1. Bone marrow regeneration after transplantation**

A. Cell viability determined by flow cytometry and expressed as percentage of total Dapi- alive cells in the BM graft at steady state, 1, 3, 8, 15 days and 3 months after bone transplantation (n=3-4). One-way ANOVA with Dunnett's multiple comparisons test used to determine statistical significance.
B. Representative whole-mount confocal z-stack projections of host and graft femurs 1 and 4 months after transplantation. Adipocytes are stained using an anti-perilipin antibody. Scale bar = 50μm. Three independent experiments yielded similar results.
C. Comparison of different hematopoietic progenitors' populations between graft and host 5 months after bon transplantation (n=4 per group).
D. Comparison of different immune cells populations between graft and host 5 months after bone transplantation (n=4 per group).

Data represented as mean ± SEM. Unless otherwise noted, statistical significance was determined using unpaired two-tailed Student's t test. *p<0.05. ** p<0.01. *** p<0.001. ****p<0.0001.

Supplemental Figure 2

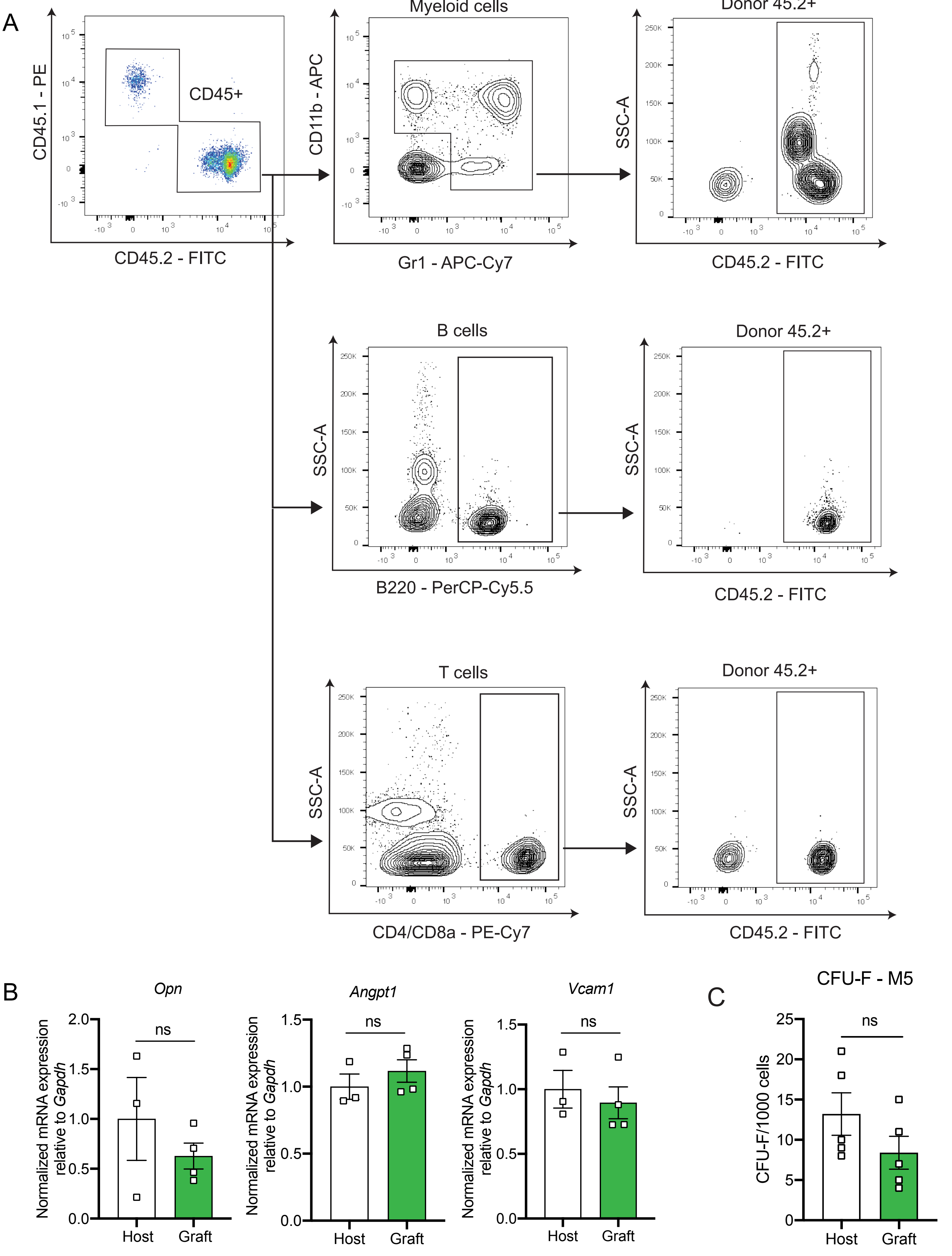

**Supplemental Figure 2. Regenerated bone marrow is capable of supporting hematopoietic cells after transplantation.**

A. Representative FACS plots showing gating strategy for trilineage reconstitution after bone marrow transplantation.
B. Quantitative RT-PCR analysis of mRNA expression of *Opn*, *Angpt1*, and *Vcam1* relative to *Gapdh* in host and graft *Nes*-GFP$^{+}$ BM-MSCs 5 months after transplantation (n=3-4 mice).
C. CFU-F absolute number of flushed CD45$^{-}$Ter119$^{-}$*Nes*-GFP$^{+}$ BM-MSCs sorted from host and graft femurs 5 months after transplantation and plated at equal number and clonal densities under CFU-F culture condition (n=5 mice per group).

Data represented as mean ± SEM. Unless otherwise noted, statistical significance was determined using unpaired two-tailed Student's t test. *p<0.05. ** p<0.01. *** p<0.001. ****p<0.0001.

Supplemental Figure 3

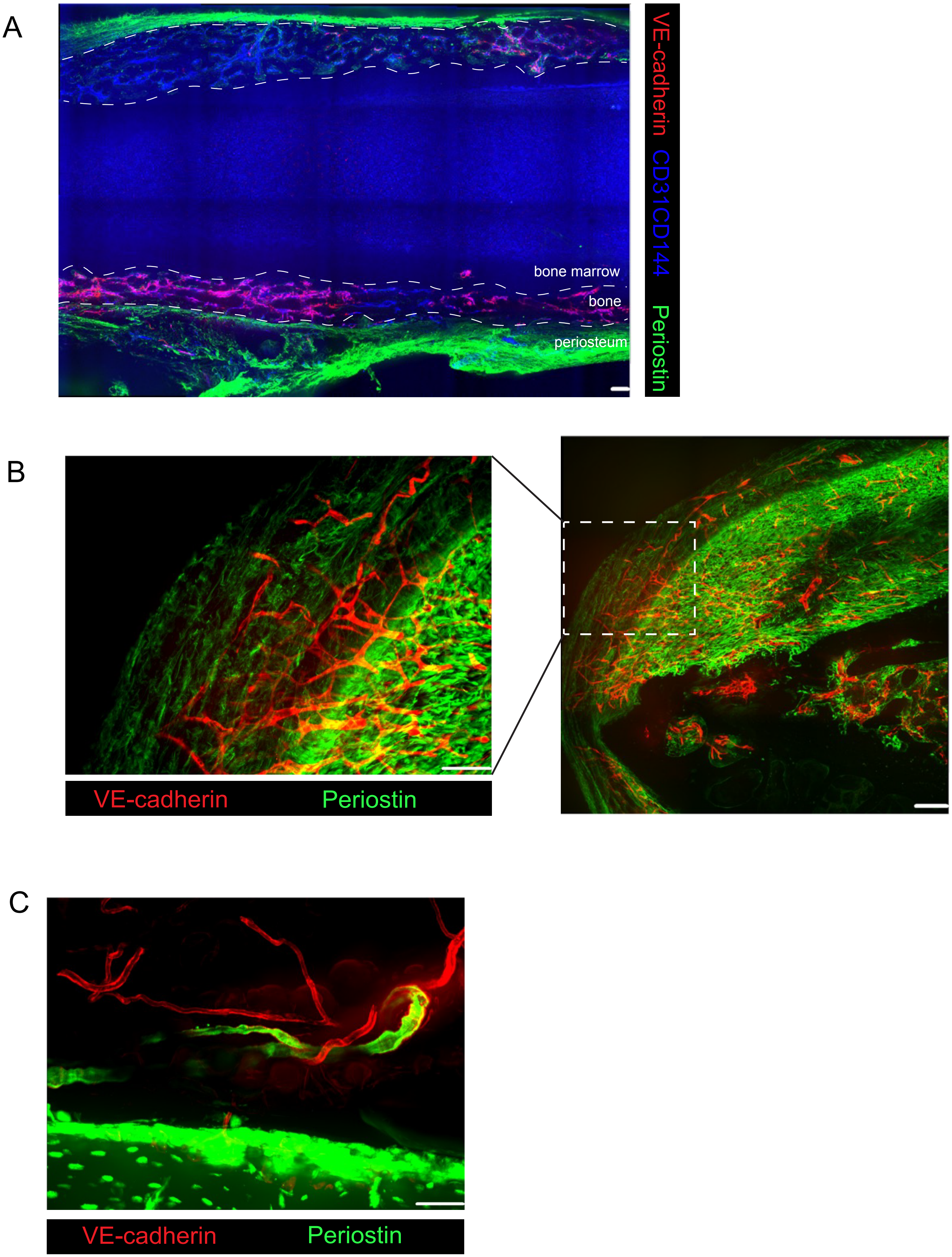

**Supplemental Figure 3. Endothelial regeneration after bone transplantation.**

A. Representative confocal image of graft VE-cadherin-cre;tdTomato femur transplanted into WT recipient fifteen days after transplantation. Extracellular matrix is labeled by an anti-periostin antibody. Scale bar = 100 μm
B. Representative confocal image of graft VE-cadherin-cre;tdTomato femur transplanted into WT recipient one month after transplantation. Extracellular matrix is labeled by an anti-periostin antibody. Scale bar = 50μm (left) and 100μm (right).
C. Representative confocal image of graft UBC-GFP femur transplanted into VE-cadherin-cre;tdTomato recipient five months after transplantation. Extracellular matrix is labeled by an anti-periostin antibody. Scale bar = 50μm.

### Supplemental Figure 4

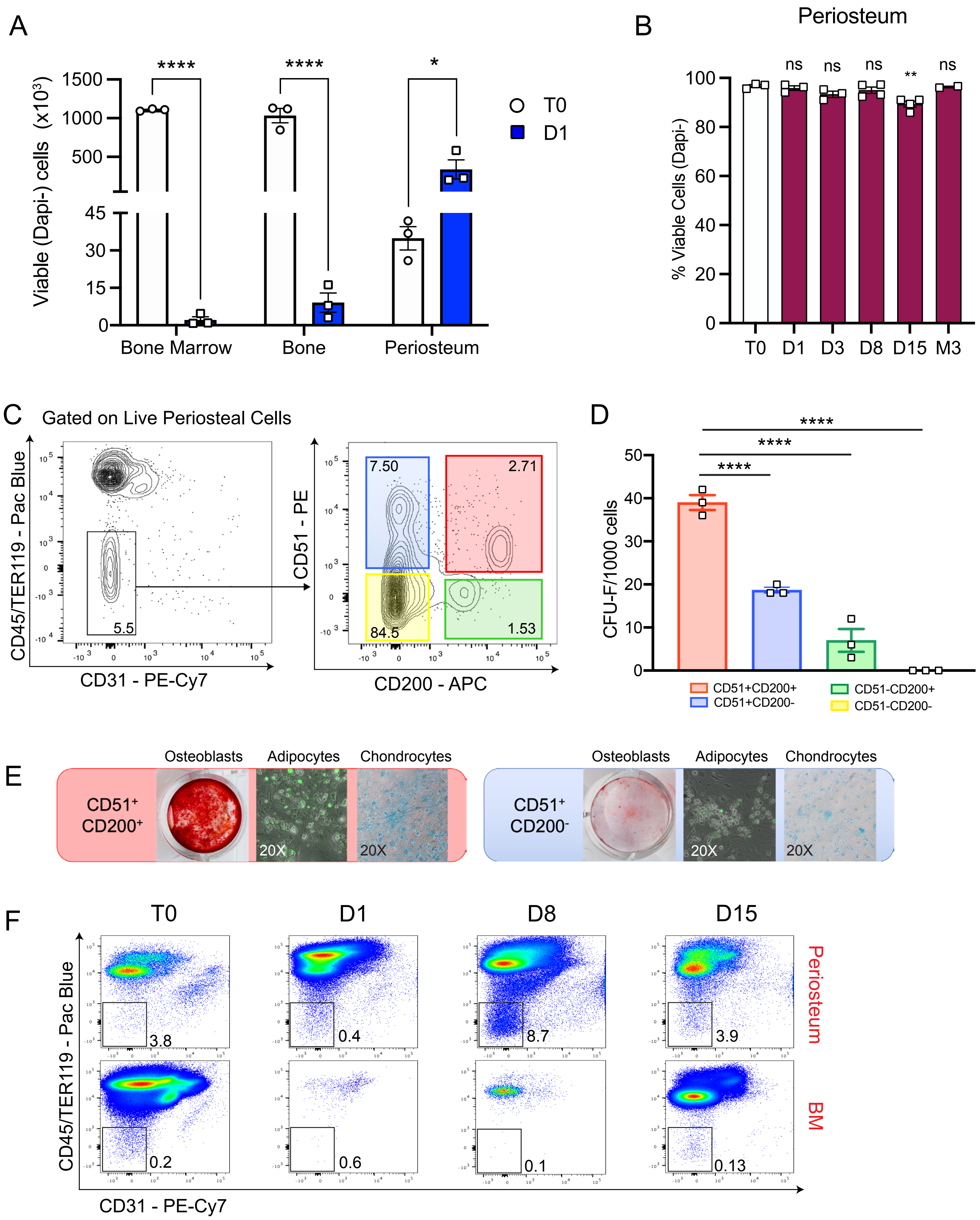

**Supplemental Figure 4. Periosteal stem cells expansion after transplantation.**

A. Number of viable cells within bone marrow, bone and periosteum at steady state and 24 hours post transplantation (n=3 per group).
B. Cell viability determined by flow cytometry and expressed as percentage of total Dapi- alive cells in the periosteum graft at steady state, 1, 3, 8, 15 days and 3 months after bone transplantation (n=2-4 per group). One-way ANOVA with Dunnett's multiple comparisons test used to determine statistical significance.
C. Representative FACS plot showing gating of $CD51^+CD200^+$ P-SSCs after gating on triple negative cells (TNC: $CD45^-Ter119^-CD31^-$) fraction.
D. Absolute CFU-F number from 1000 sorted cells within various TNC subsets of the periosteum. One-way ANOVA with Dunnett's multiple comparisons test used to determine statistical significance.
E. Representative trilineage differentiation pictures of $CD51^+CD200^+$ compared to the group with the next highest CFU-F activity ($CD51^+CD200^-$ cells). Differentiation into osteoblasts, adipocytes and chondrocytes observed using Alizarin Red, Bodipy, and Alican Blue staining, respectively.
F. Representative FACS plots of graft periosteum and bone marrow analysis gated on TNC fraction at steady state, one-, eight- and fifteen-days post transplantation.

Data represented as mean ± SEM. Unless otherwise noted, statistical significance was determined using unpaired two-tailed Student's t test. *p<0.05. ** p<0.01. *** p<0.001. ****p<0.0001.

### Supplemental Figure 5

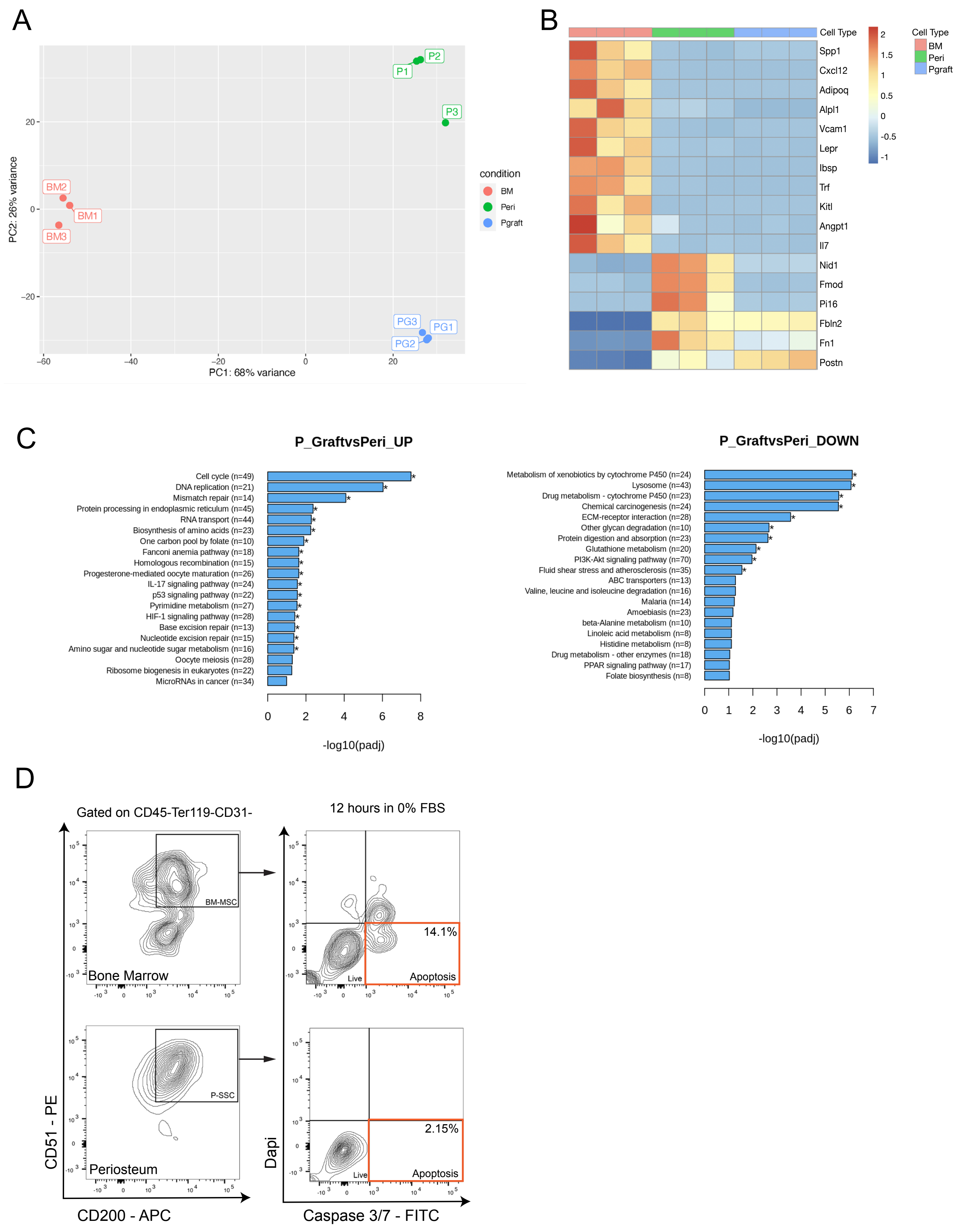

**Supplemental Figure 5. RNA sequencing analysis of periosteal SSCs, three-day graft periosteal and SSCs, and bone marrow MSCs.**

A. 2D principal component analysis (PCA) plot showing variance between steady-state P-SSCs and graft P-SSCs three days after bone transplantation (n=3).
B. Heatmap expression level of selected genes as defined by previous studies for HSC nice cell and extracellular matrix genes (n=3).
C. KEGG analysis displaying signatures that are significantly up- and down-regulated between steady-state P-SSCs and graft P-SSCs three days after bone transplantation (n=3).
D. Representative FACS plots showing gating strategy for analyzing apoptosis 12 hours after switching from 20% to 0% FBS media

Data represented as mean ± SEM. Unless otherwise noted, statistical significance was determined using unpaired two-tailed Student’s t test. *$p<0.05$. ** $p<0.01$. *** $p<0.001$. ****$p<0.0001$.

#### Supplemental Figure 6

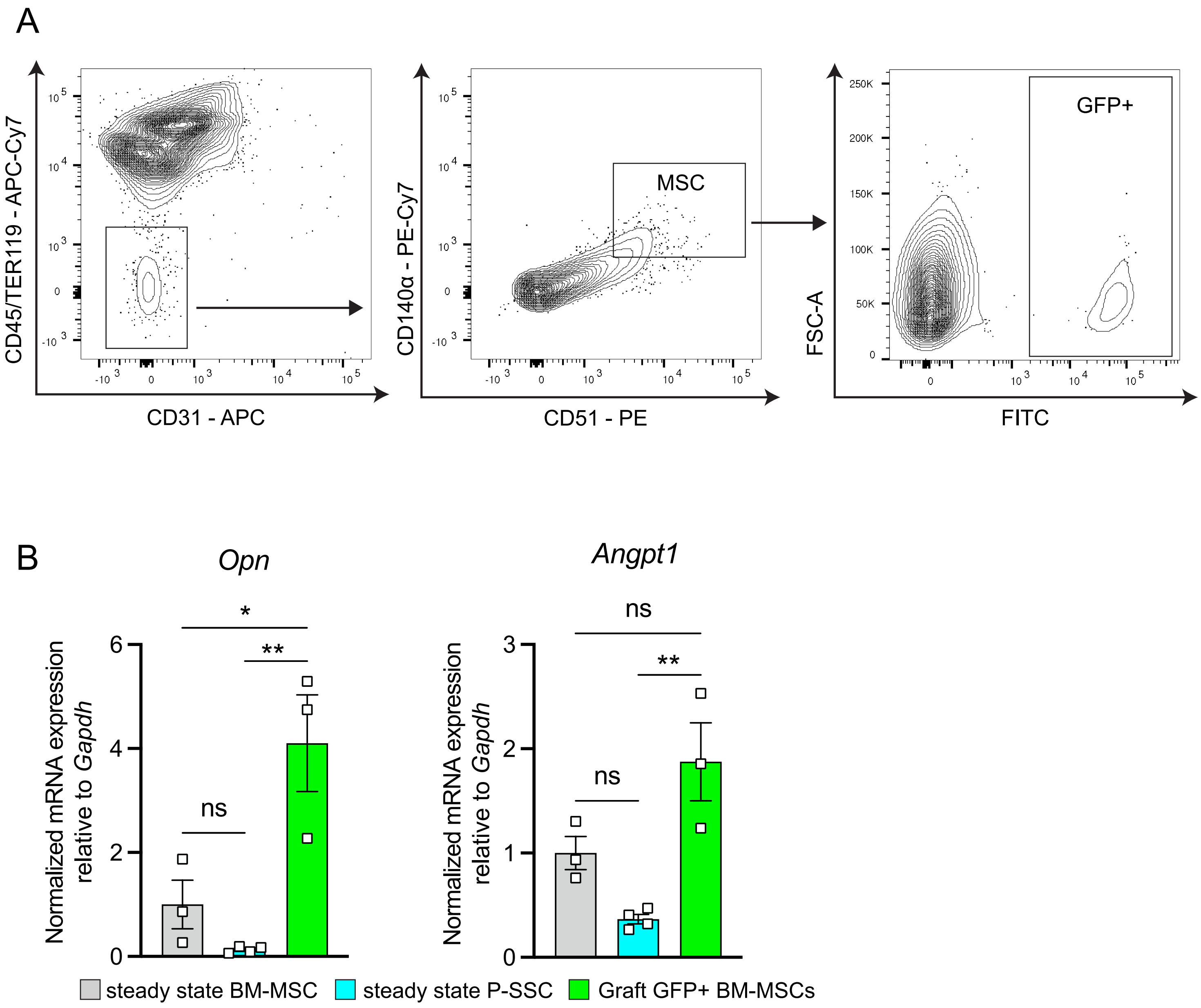

**Supplemental Figure 6. Periosteal SSCs migrate into the bone marrow and upregulates HSC-niche factors.**

A. Representative FACs plots showing gating strategy for sorting GFP+ BM-MSCs used in qPCR analysis.
B. Quantification of *Opn* and *Angpt1* mRNA level relative to *Gapdh* in sorted control bone marrow $CD45^-Ter119^-CD31^-Nestin\text{-}GFP^+$ BM-MSCs, $CD45^-Ter119^-CD31^-CD51^+CD200^+$ steady-state P-SSCs, and $CD45^-Ter119^-CD31^-CD51^+CD200^+GFP^+$ graft BM-MSCs (n=3-4 per group). One-way ANOVA used to determine statistical significance.

Data represented as mean ± SEM. Unless otherwise noted, statistical significance was determined using unpaired two-tailed Student's t test. *p<0.05. ** p<0.01. *** p<0.001. ****p<0.0001.

Supplemental Figure 7

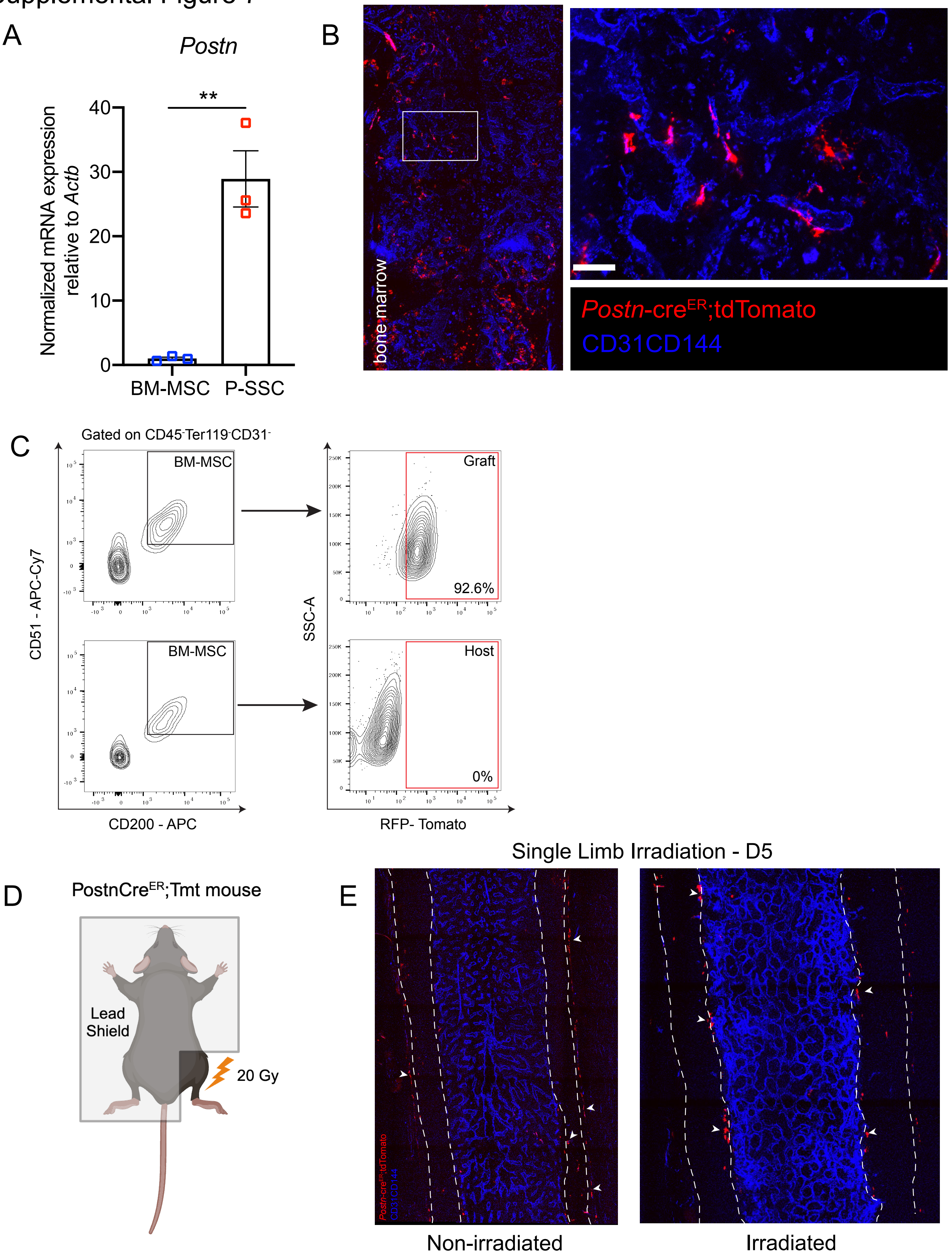

**Supplemental Figure 7. *Postn*-Cre$^{ER;}$tdTomato can be used to lineage trace periosteal SSCs and label periosteum-derived BM-MSCs.**

A. Postn expression in sorted, steady-state CD51+CD200+ BM-MSCs and P-SSCs from CD45.2 WT mice (n=3).
B. Representative whole-mount confocal z-stack projections of bone marrow from *Postn*-Cre$^{ER}$;tdTomato femurs into a WT recipient mouse 5 months after transplantation. Three independent experiments yielded similar results. Scale bar = 50μm (right panel)
C. Representative FACS plots showing gating strategy for identifying Tomato$^{+}$ BM-MSCs
D. Schematic diagram representing single limb irradiation of *Postn*-Cre$^{ER}$;tdTomato mice
E. Whole-mount confocal z-stack projections of bone marrow 5 days later. White arrows point at areas of Tomato labeling along the periosteum and cortical bone.

Data represented as mean ± SEM. Unless otherwise noted, statistical significance was determined using unpaired two-tailed Student’s t test. *p<0.05. ** p<0.01. *** p<0.001. ****p<0.0001.

Supplemental Figure 8

A

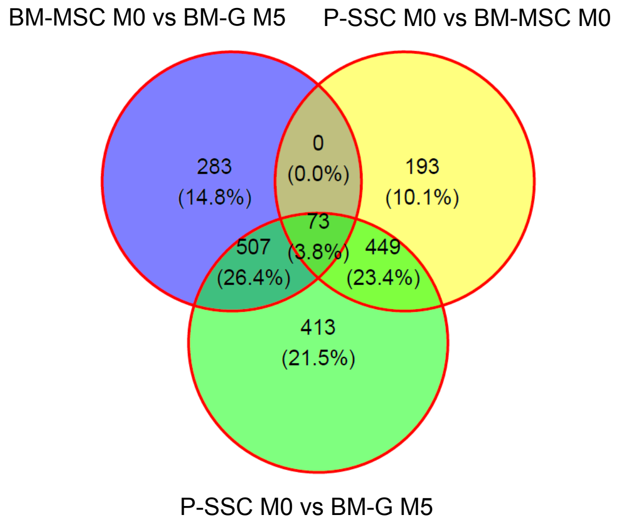


B

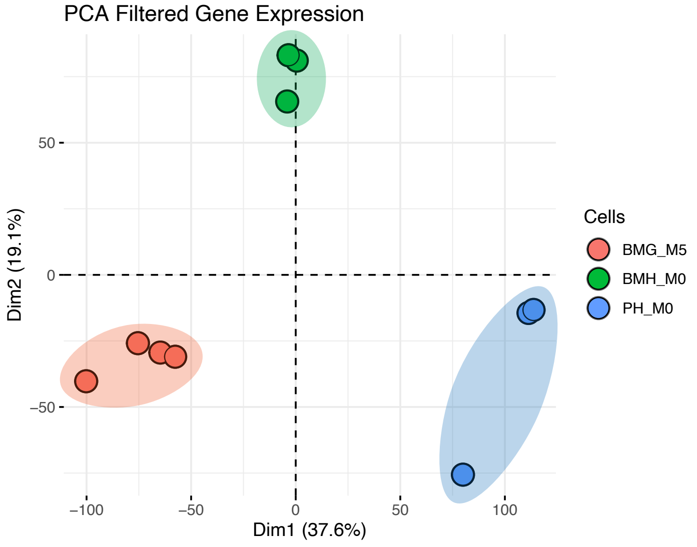

**Supplemental Figure 8: Graft Tomato+ BM-MSCs derived from P-SSCs exhibit profile similar to that of steady-state BM-MSCs.**

A. Venn diagram of RNA sequencing analysis comparing the numbers of differentially expressed genes between the groups.
B. 2D principal component analysis (PCA) plot showing variance between graft BM-MSCs at five months post-transplantation, and BM-MSCs and P-SSCs at steady state (n=3-4).

Data represented as mean ± SEM. Unless otherwise noted, statistical significance was determined using unpaired two-tailed Student's t test. *$p<0.05$. ** $p<0.01$. *** $p<0.001$. ****$p<0.0001$.
